# cMPL-Based Purification and Depletion of Human Hematopoietic Stem Cells: Implications for Pre-Transplant Conditioning

**DOI:** 10.1101/2024.02.24.581887

**Authors:** Daisuke Araki, Sogun Hong, Nathaniel Linde, Bryan Fisk, Neelam Redekar, Christi Salisbury-Ruf, Allen Krouse, Theresa Engels, Justin Golomb, Pradeep Dagur, Diogo M. Magnani, Zhirui Wang, Andre Larochelle

## Abstract

The transplantation of gene-modified autologous hematopoietic stem and progenitor cells (HSPCs) offers a promising therapeutic approach for hematological and immunological disorders. However, this strategy is often limited by the toxicities associated with traditional conditioning regimens. Antibody-based conditioning strategies targeting cKIT and CD45 antigens have shown potential in mitigating these toxicities, but their long-term safety and efficacy in clinical settings require further validation. In this study, we investigate the thrombopoietin (TPO) receptor, cMPL, as a novel target for conditioning protocols. We demonstrate that high surface expression of cMPL is a hallmark feature of long-term repopulating hematopoietic stem cells (LT-HSCs) within the adult human CD34+ HSPC subset. Targeting the cMPL receptor facilitates the separation of human LT-HSCs from mature progenitors, a delineation not achievable with cKIT. Leveraging this finding, we developed a cMPL-targeting immunotoxin, demonstrating its ability to selectively deplete host cMPL^high^ LT-HSCs with a favorable safety profile and rapid clearance within 24 hours post-infusion in rhesus macaques. These findings present significant potential to advance our understanding of human hematopoiesis and enhance the therapeutic outcomes of *ex vivo* autologous HSPC gene therapies.

Graphical abstract

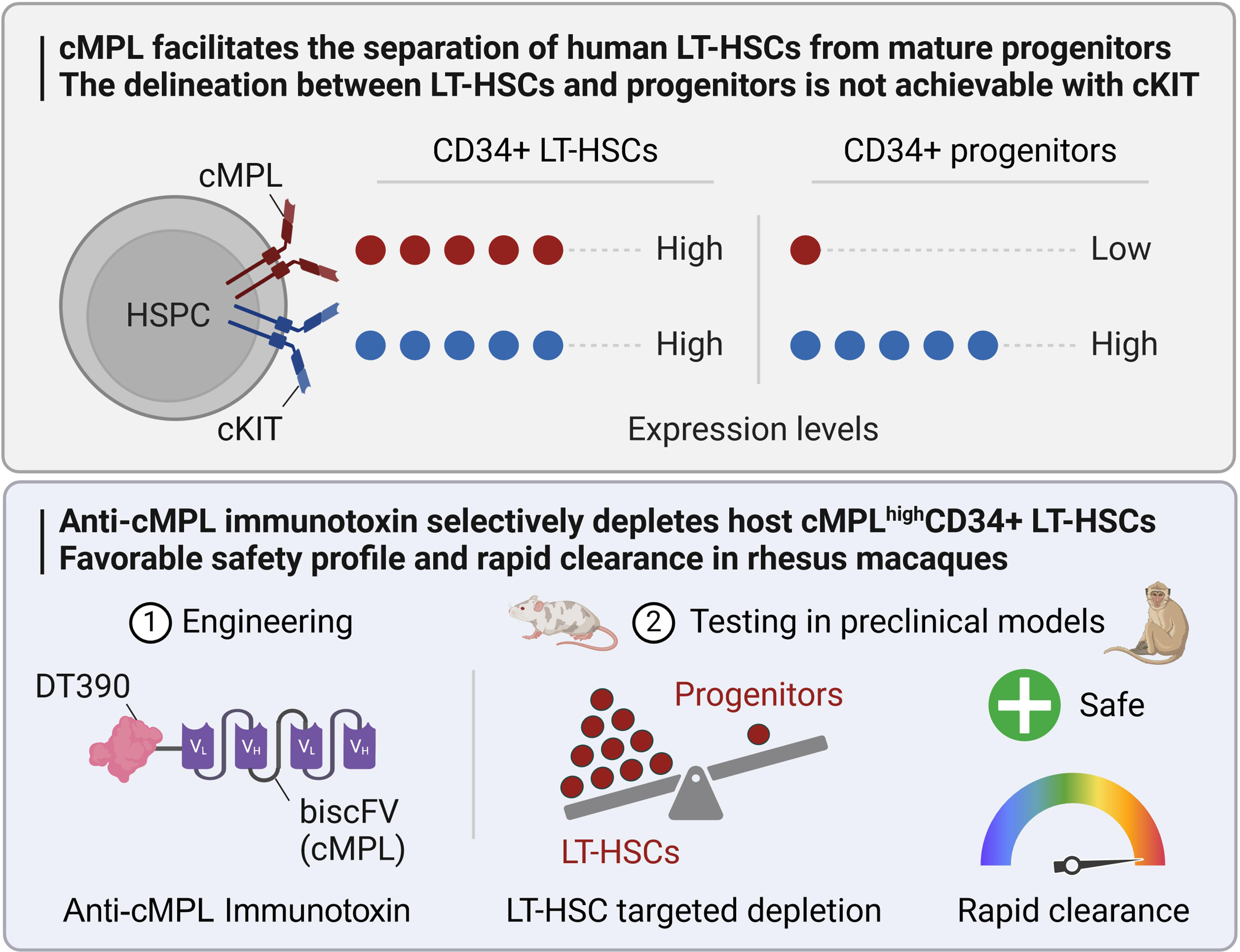

## Introduction

*Ex vivo* autologous hematopoietic stem and progenitor cell (HSPC) gene therapy has emerged as a transformative treatment for inherited blood disorders^1–3^. The success of this therapy hinges on the conditioning regimen to prepare a conducive niche within the recipient’s bone marrow (BM) for the engraftment of infused genetically modified HSPCs^4^. Traditional conditioning methods have predominantly utilized non-myeloablative doses of busulfan, either in isolation or in conjunction with other cytotoxic agents^4–9^. However, these regimens are associated with a spectrum of acute and long-term toxicities, such as mucositis, infections, endocrine dysfunctions, and secondary malignancies^5,10–12^. Thus, alternative strategies to promote engraftment of genetically modified HSPCs with increased safety warrant investigation. Recent preclinical investigations have demonstrated promising advances in the depletion of HSPCs using monoclonal antibodies, potentially offering a safer and more effective alternative to traditional conditioning regimens^13–19^. Key studies have focused on antibodies targeting human c-KIT (CD117), with promising results in HSPC depletion in murine^13,15,16^ and non-human primate (NHP) models^20,21^. Parallel investigations involving immunotoxin-conjugated monoclonal antibodies targeting CD45 have also shown potential in murine models^19,22,23^. However, the clinical application of these novel strategies necessitates thorough evaluation of long-term safety and efficacy in humans. The broad expression of these antigens across various cell types and tissues raises concerns about off-target effects. CD45 expression is ubiquitous in hematopoietic cells, while cKIT is broadly expressed across both hematopoietic stem cells (HSCs) and progenitors, as well as germ cells, melanocytes, mast cells, cardiac progenitors, cajal cells of the gastrointestinal tract, epithelial cells in skin adnexa, breast, and subsets of cerebellar neurons^24^. The use of anti-cKIT and anti-CD45-based conditioning regimens can result in significant cytopenias, leading to prolonged hospitalizations, increased infection risks, and higher transfusion needs, thus exacerbating patient morbidity and healthcare costs. Therefore, particularly in contexts like *ex vivo* autologous HSPC gene therapy where preservation of immunity is desired, there is a pressing need for a non-genotoxic conditioning approach that more precisely targets long-term repopulating HSCs (LT-HSCs) to address these challenges and facilitate broader clinical implementation.

The hematopoietic cytokine thrombopoietin (TPO) and its cellular receptor cMPL (CD110) have recently garnered attention beyond their established function in megakaryopoiesis^25,26^. Recent insights reveal TPO as a key regulator of HSC quiescence, survival, and proliferation, modulating critical cell signaling pathways upon binding to cMPL^27–31^. The effectiveness of eltrombopag, a TPO receptor agonist, in restoring trilineage hematopoiesis in patients with immune aplastic anemia^32–35^, underscores the pivotal role of the TPO:cMPL axis in HSC regulation. Importantly, cMPL expression is largely restricted to megakaryopoiesis and early hematopoietic stages, with minimal expression in mature blood cells or non-hematopoietic tissues^36^.

Here, we have engineered a truncated diphtheria toxin-based recombinant anti-cMPL immunotoxin and explored its potential as a novel and safer conditioning alternative in HSPC transplantation. Our research shows that high surface expression of cMPL is a reliable marker for LT-HSCs within the adult human CD34+ cell population. Furthermore, our pre-clinical investigations in xenograft murine models and NHPs reveal that cMPL-specific immunotoxins can selectively and safely eradicate primitive cMPL^high^ LT-HSCs. These findings pave the way for an anti-cMPL-based non-genotoxic conditioning approach, offering a promising strategy to optimize outcomes in *ex vivo* autologous HSPC gene therapies.

## Results

### High cMPL surface expression enriches phenotypically and transcriptionally defined human LT-HSCs

Human HSPC activity is regulated by the TPO:cMPL and stem cell factor (SCF):cKIT signaling axes. The expression profile of the cKIT receptor has been extensively studied within murine HSPCs^37–39^, and antibodies targeting cKIT have subsequently been leveraged in pre-clinical studies as a novel conditioning regimen^20,21^. The cMPL receptor is also known to be highly expressed in quiescent murine LT-HSCs^27,28^. However, the expression profiles of these receptors in adult human HSPCs have not been comprehensively characterized. We conducted a detailed flow cytometry analysis to delineate the surface expression patterns of cMPL and cKIT receptors on G-CSF mobilized peripheral blood (MPB) CD34+ HSPCs derived from healthy donors. This analysis employed a panel of established human LT-HSC surface markers (CD34, CD38, CD90, CD45RA and CD49f) and antibodies specific for cMPL and cKIT (Figure 1A). We found that both cMPL and cKIT receptors were expressed in over 95% of the CD34+ cell population derived from MPB. Notably, cMPL demonstrated a higher expression level within the subset enriched with LT-HSCs, distinctly marked by CD34+CD38-CD45RA-CD90+CD49f+ phenotype, in comparison to the CD34+CD38+ hematopoietic progenitor cell subset and the more broadly defined CD34+ cell population (Figures 1A, B). Conversely, the expression levels of cKIT did not differentiate between LT-HSCs and other subsets, including both the CD34+CD38+ cell population and the bulk CD34+ cell pool (Figures 1A, C).

**Figure 1.**
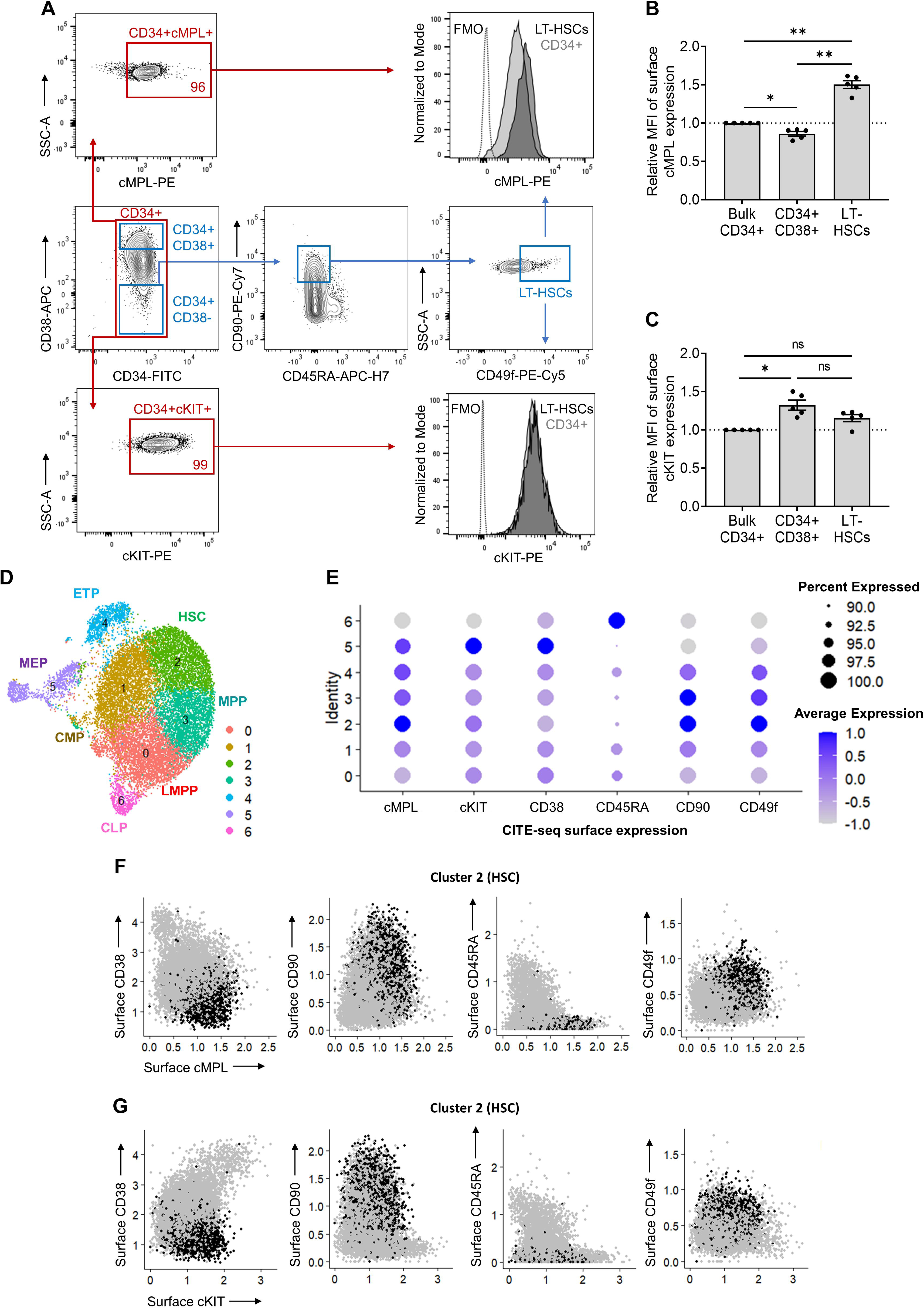
High surface expression of cMPL enriches phenotypically and transcriptionally defined human HSCs. (A) Flow cytometry gating strategy employed to resolve distinct populations within the CD34+ cell fraction of human mobilized peripheral blood (MPB). Initial gating on CD34+ cells (red gate) allowed for the subsequent separation into progenitors (CD34+CD38+) and LT-HSCs (CD34+CD38-CD90+CD45RA-CD49f+) (blue gates). The expression profiles of cMPL and cKit receptors were then evaluated within these subsets. The histograms on the right display representative mean fluorescence intensity (MFI) for cMPL and cKIT, illustrating their expression levels within the LT-HSC and CD34+ cell populations. Fluorescence minus one (FMO) is included as a control. (B, C) Quantitation of cMPL and cKIT surface expression in identified populations by MFI, normalized against the expression within total CD34+ cells (n = 5 independent donors). (D) Uniform Manifold Approximation and Projection (UMAP) clustering from CITE-seq data of human MPB CD34+ cells, identifying seven distinct HSPC clusters: lymphoid-primed multipotent progenitors (LMPP, cluster 0), common myeloid progenitors (CMP, cluster 1), hematopoietic stem cells (HSC, cluster 2), multipotent progenitors (MPP, cluster 3), early T-lineage progenitors (ETP, cluster 4), megakaryocytic-erythroid progenitors (MEP, cluster 5) and common lymphoid progenitors (CLP, cluster 6). (E) Dot plot visualization of protein surface expression from CITE-seq, with dot size reflecting cell expression percentage and color intensity denoting scaled average expression level across cluster types. (F, G) Feature plots from Seurat analysis correlating cMPL (F) and cKIT (G) expression with established HSC phenotypes within the transcriptionally defined HSC cluster 2. Datapoints in panels (B) and (C) represent the mean ± standard error of the mean (SEM), with statistical significance assessed via one-way ANOVA with Tukey correction. Notations of statistical significance are as follows: ns; not significant, * p ≤ 0.05, ** p ≤ 0.01.

To corroborate these observations at the transcriptional level, we performed cellular indexing of transcriptomes and epitopes by sequencing (CITE-seq) on MPB CD34+ cells from a healthy donor. The CITE-seq approach incorporated targeted antibodies against cMPL and cKIT surface receptors, along with the aforementioned HSC epitopes. After quality control, 16,767 single CD34+ cells were analyzed. Summary statistics are provided in Supplemental Figure 1A. Uniform manifold approximation and projection (UMAP) analysis delineated seven distinct HSPC clusters (Figure 1D), confirmed via HSPC surface markers (Figure 1E) and lineage marker gene expression (Supplemental Figure 1B). The transcriptionally defined HSC cluster (cluster 2) displayed the highest cMPL expression (Figure 1E), aligning with our phenotypic flow cytometry data (Figures 1A-C). Furthermore, cells with higher cMPL expression consistently mirrored the established LT-HSC immunophenotype, marked by reduced CD38 and CD45RA expression and elevated CD90 and CD49f levels, reinforcing the robust association between high cMPL expression and the prototypical LT-HSC profile (Figure 1F). In contrast, CD34+ cells with higher cKIT receptor levels were not enriched in HSC cluster 2 population (Figures 1E, G), but were more prominent in the MEP cluster (cluster 5), with indistinct levels among HSCs and other progenitor populations (Figure 1E). These results collectively suggest that high surface expression of the cMPL receptor serves as a defining characteristic of phenotypically and transcriptionally defined adult human LT-HSCs, providing a more precise and selective LT-HSC marker compared to cKIT expression.

### High cMPL expression on human CD34+ cells correlates with robust long-term repopulating capacity *in vivo*

To investigate the potential of cMPL surface expression as a marker for isolating human HSCs with long-term engraftment capacity *in vivo*, we conducted a xenotransplantation assay. Human MPB CD34+ cells were segregated into cMPL^high^ (top 10%) and cMPL^low^ (bottom 10%) populations via fluorescence-activated cell sorting (FACS). We then transplanted 5 x 10^4^ cells from each subset into immunodeficient NOD,B6.SCID IL-2r ^-^/^-^Kit^W41/W41^ (NBSGW) mice. Additionally, to benchmark engraftment efficiency, we transplanted a fivefold larger number of cMPL^low^CD34+ cells (2.5 x 10^5^). Concurrently, the reliability of cKIT surface expression in identifying adult LT-HSCs was assessed as a comparative measure (Figure 2A).

**Figure 2.**
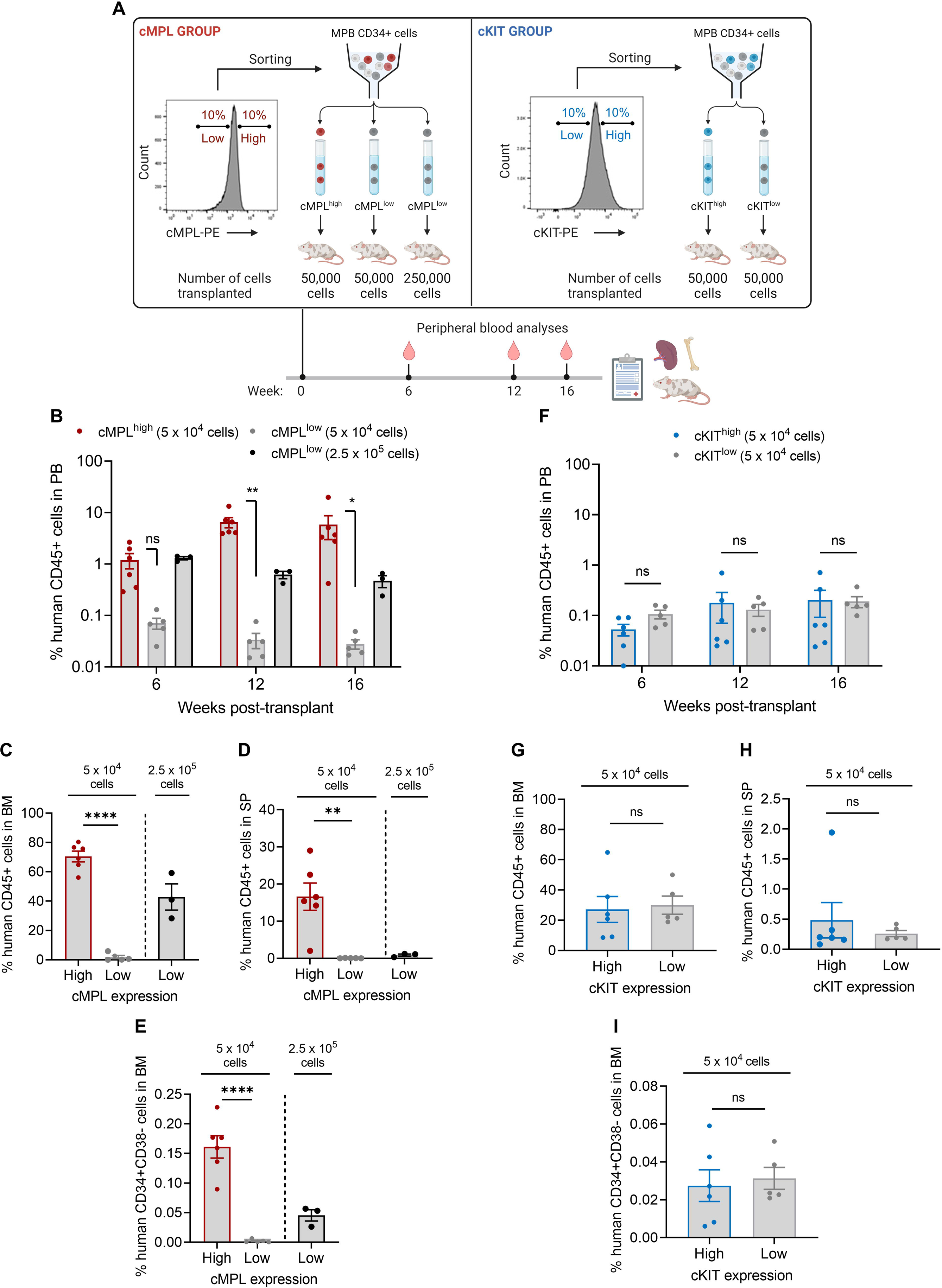
The cMPL receptor serves as a marker for human long-term repopulating HSCs. (A) Schematic of the experimental procedure: Human CD34+ cells from mobilized peripheral blood (MPB) were sorted into high and low subsets based on surface expression of cMPL and cKIT, followed by transplantation into NBSGW mice to assess hematopoietic reconstitution (n = 3-5 mice per group). (B-E) Comparative analysis of human cell engraftment in the peripheral blood (PB) (B), bone marrow (BM) (C), and spleen (SP) (D) of NBSGW mice after transplantation with human cMPL^high^CD34+ or cMPL^low^CD34+ cells, with panel (E) depicting the engraftment of HSC-enriched populations (CD34+CD38-) within the BM. (F-I) Comparative analysis of human cell engraftment in the PB (F), BM (G), and SP (H) of NBSGW mice after transplantation with human cKIT^high^ or cKIT^low^ CD34+ cells, with panel (I) depicting the engraftment of HSC-enriched populations (CD34+CD38-) within the BM. Datapoints represent the mean ± standard error of the mean (SEM), with statistical significance assessed via two-sided unpaired t-tests. Notations of statistical significance are as follows: ns, not significant, * p ≤ 0.05, ** p ≤ 0.01, **** p ≤ 0.0001.

During the 16-week observation period, serial peripheral blood (PB) sampling from the primary recipient mice revealed a stark contrast in hematopoietic reconstitution between the cMPL^high^ and cMPL^low^ groups (Figure 2B). Mice infused with cMPL^high^CD34+ cells showed a progressive increase in human cell chimerism. Conversely, mice receiving cMPL^low^CD34+ cells displayed a waning pattern of engraftment. Notably, by the 16^th^ week post-transplant, chimerism in the PB, BM, and spleen (SP) was significantly higher in the cMPL^high^ group, with increases of 209, 37, and 283-fold, respectively, compared to the cMPL^low^ group (Figures 2B-D). Although the lineage distribution within human CD45+ cell compartments across PB, BM and SP was comparable between the cMPL^high^ cMPL^low^ groups at the study endpoint (Supplemental Figure 2A), the frequency of the LT-HSC-enriched CD34+CD38-cell fraction in the BM of transplanted mice was notably 72-fold higher in the cMPL^high^ group compared to the cMPL^low^ group (Figure 2E). In contrast, chimerism levels in the PB, BM, and SP of mice transplanted with cKIT^high^ and cKIT^low^ cells demonstrated no significant differences, highlighting a marked difference between HSPCs enriched for the cKIT and cMPL receptors (Figures 2F-I, Supplemental Figure 2B).

To further substantiate the selective enrichment of LT-HSCs based on cMPL expression levels, we performed a secondary transplantation assay using a limiting dilution approach. Human CD45+ cells isolated from the BM of primary recipient mice from both cMPL^high^ and cMPL^low^ groups were transplanted into secondary NBSGW mice. For the cMPL^low^ group, assessments were limited to recipient mice initially transplanted with 2.5 x 10^5^ cells due to insufficient engraftment in animals transplanted with the lower cell dose. The BM engraftment was evaluated after 16 weeks, cumulating a total engraftment period of 32 weeks (Figure 3A). A significantly higher proportion of secondary recipient mice exhibited BM engraftment when transplanted with cells from the cMPL^high^ primary group compared to the cMPL^low^ group (37% versus 6%, respectively) (Figure 3B). Extreme limiting dilution analysis revealed a 7.9-fold greater frequency of self-renewing LT-HSCs within the human CD45+ cells from the cMPL^high^ group compared to the cMPL^low^ group (p=0.009) (Figure 3C). To account for initial differences in the number of human CD34+ cells transplanted into primary recipients and the resulting human BM chimerism between the cMPL^high^ and cMPL^low^ groups, we quantified absolute LT-HSC numbers harvested from four hindlimb and two pelvic bones of primary mice. This quantitation facilitated the calculation of LT-HSC frequencies within the original human cMPL^high^ or cMPL^low^ HSPC populations. The results revealed a substantial 152-fold enrichment of LT-HSCs within the cMPL^high^CD34+ cell fraction, with a frequency of 1 LT-HSC per 1,266 cells, compared to a sparse frequency of 1 per 192,308 cells within the cMPL^low^CD34+ cell fraction (Figure 3D). Collectively, our findings provide compelling evidence that high cMPL surface expression is a dependable marker to enrich LT-HSCs within the adult human CD34+ cell pool, whereas cKIT surface expression does not offer a similarly effective distinction between LT-HSCs and their more differentiated progeny within the hematopoietic hierarchy.

**Figure 3.**
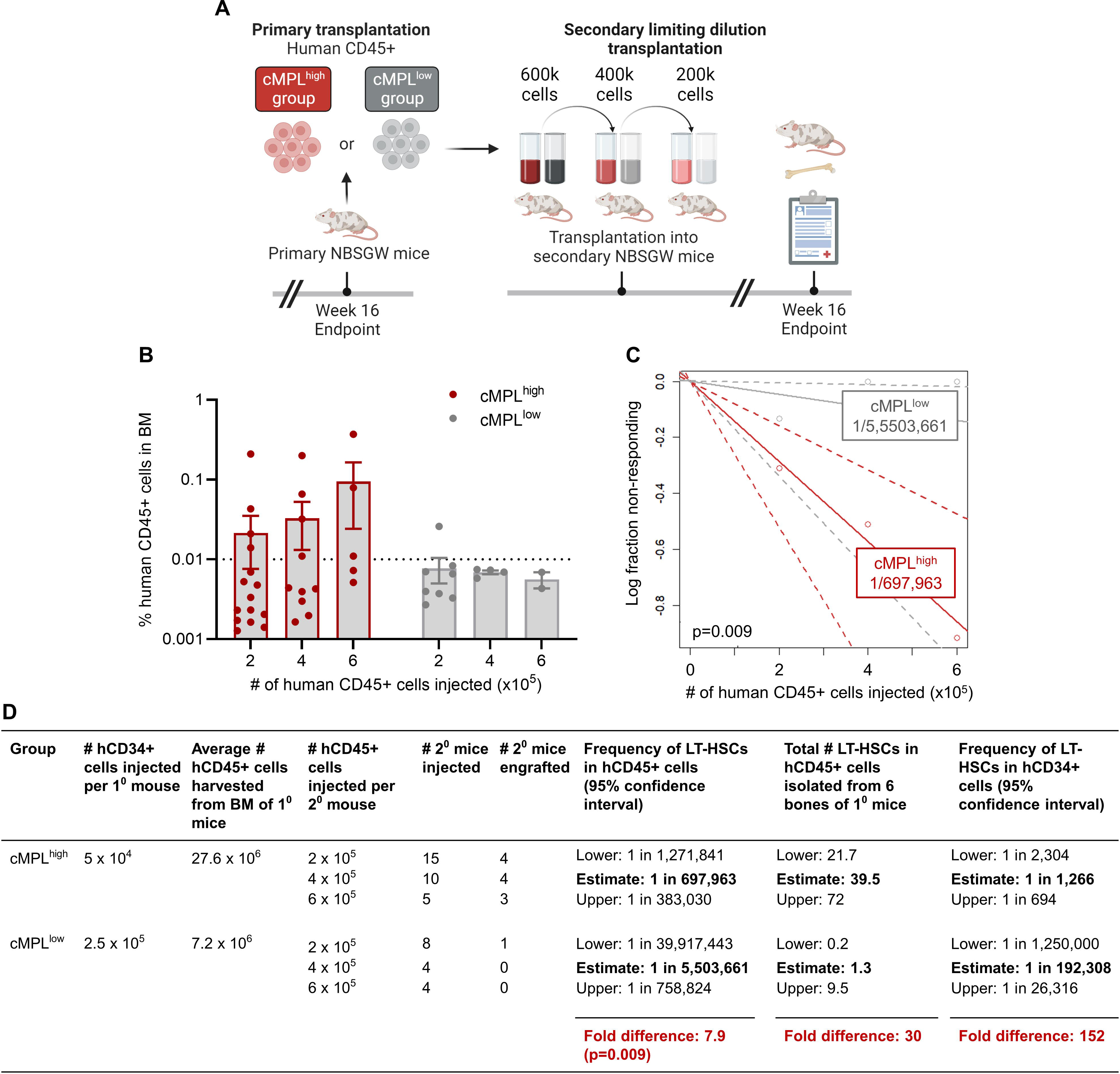
Significant enrichment of human LT-HSCs within cMPL^high^CD34+ cells measured by a limiting-dilution secondary transplantation assay. (A) Schematic of the experimental procedure: Human CD45+ cells, harvested from the bone marrow (BM) of primary recipient mice from both cMPL^high^ and cMPL^low^ groups, underwent a secondary transplantation into NBSGW immunodeficient mice. The cells were transplanted at a limiting dilution after conditioning with a low busulfan dose, and BM engraftment levels were assessed 16 weeks following the transplantation (n = 4-15 mice per group). (B) Quantitative analysis of human CD45+ cell populations within the BM after secondary transplantation of cells derived from both cMPL^high^ and cMPL^low^ groups. The dashed line represents the threshold (0.01%) above which the presence of human CD45+ cells, encompassing both myeloid (CD13+) and lymphoid (CD20+) lineages, is considered indicative of successful secondary engraftment. (C) Semilogarithmic representation of LT-HSC frequencies within the human CD45+ population post-secondary transplant, comparing the cMPL^high^ and cMPL^low^ groups. The solid lines represent the best-fit linear regression for each group, while dotted lines indicate the 95% confidence intervals. (D) Quantitation of LT-HSC frequencies and absolute numbers derived from secondary transplantation assays conducted at limiting dilutions. LT-HSC counts within hCD45+ cells (column 8) were determined by multiplying LT-HSC frequencies (column 7) by the average pooled numbers of human CD45+ cells harvested from the primary mice’s limbs, including 4 hindlimb and 2 pelvic bones. Frequencies of LT-HSCs within human CD34+ cells (column 9) were established by dividing the LT-HSC numbers (column 8) by the initial quantity of hCD34+ cells transplanted into each primary mouse (column 2). The associated p-value was calculated using extreme limiting dilution statistics. Datapoints in panel (B), represent the mean ± standard error of the mean (SEM).

### DT390-biscFV(cMPL) immunotoxin impairs the proliferation of cMPL-expressing human cell lines and HSPCs *in vitro*

The observation that cMPL is predominantly expressed in LT-HSCs rather than in more mature hematopoietic progenitors laid the groundwork for the development of a targeted conditioning approach for the selective depletion of LT-HSCs. We engineered a novel immunotoxin, DT390-biscFV(cMPL), by fusing a bivalent anti-cMPL single-chain fragment variable (biscFV(cMPL)) with diphtheria toxin (DT) truncated at amino acid residue 390 (DT390) (Figure 4A, B). The biscFV(cMPL) segment, a humanized antibody derivative, has been validated for its therapeutic efficacy in patients with thrombocytopenia and demonstrates a robust binding affinity for cMPL receptors in humans and NHPs^40^. The design of DT390 integrates the catalytic and transmembrane domains of DT but excludes the native receptor-binding domain, thereby enhancing the specific toxicity towards cMPL+ cells and minimizing non-targeted binding interactions^41–43^.

**Figure 4.**
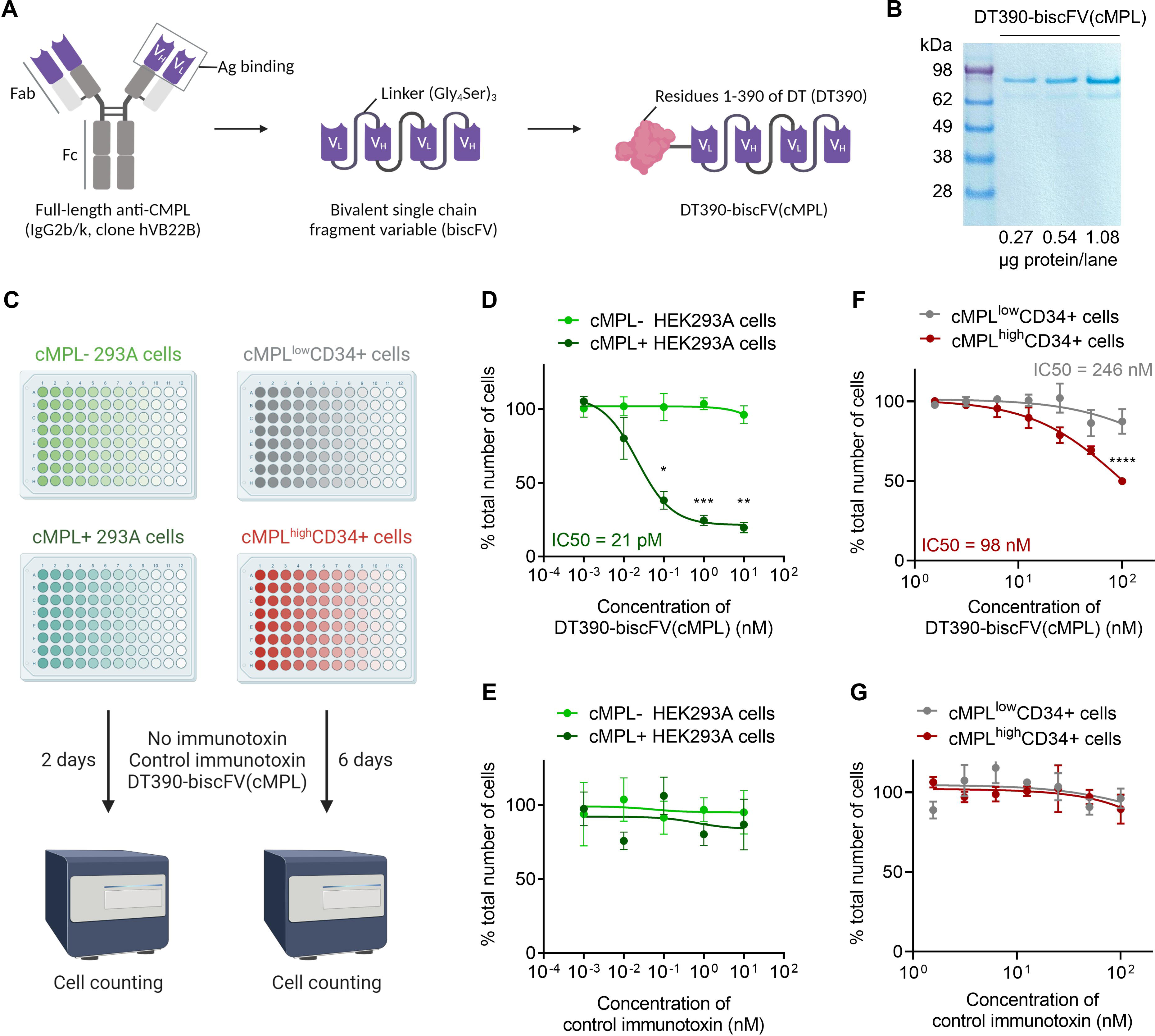
DT390-biscFV(cMPL) immunotoxin impairs the proliferation of cMPL-expressing human cell lines and HSPCs *in vitro.* (A) Diagram of DT390-biscFV(cMPL) depicting the configuration of variable light (V_L_) and variable heavy (V_H_) chain sequences from the cMPL-specific monoclonal antibody hVB22B. These sequences are joined by (Gly_4_Ser)_3_ linkers, with the truncated diphtheria toxin (DT390) attached at the N-terminus of the biscFV(cMPL) to facilitate targeted cytotoxicity. (B) SDS-PAGE analysis of DT390-biscFV(cMPL) (molecular weight = 97 kDa). Lane 1: protein marker; Lanes 2 – 4: 0.27, 0.54 and 1.08 μg of DT390-biscFV(cMPL), respectively. (C) Schematic of the experimental procedure: *In vitro* cytotoxicity assay was performed using human cMPL-expressing HEK293A cells, control (cMPL-) HEK293A cells, human cMPL^high^ mobilized peripheral blood (MPB) CD34+ cells or human cMPL^low^ MPB CD34+ cells. Cells were treated with DT390-biscFV(cMPL) or a non-specific control immunotoxin for 2 days (HEK293A cells) or 6 days (CD34+ cells). Cellular growth was assessed by automated counting, with values normalized to untreated control cultures. (D, E) Response of HEK293A cells to DT390-biscFV(cMPL) (D) or control immunotoxin (E) following a 2-day culture (n = 3 independent experiments). (F, G) Response of human MPB CD34+ cells to DT390-biscFV(cMPL) (F) or control immunotoxin (G) after a 6-day culture (n = 3 independent donors). Datapoints represent the mean ± standard error of the mean (SEM), with statistical significance assessed via two-sided unpaired t-tests. Notations of statistical significance are as follows: * p ≤ 0.05, ** p ≤ 0.01, *** p ≤ 0.001, **** p ≤ 0.0001.

To assess the specific cytotoxic effect of DT390-biscFV(cMPL) on cMPL-expressing cells, cell proliferation assays were performed on HEK293A cells, engineered to express human cMPL, after treatment with increasing concentrations of the immunotoxin. DT390-scFV(C21), targeting an unrelated antigen, served as a control to evaluate non-specific effects (Figure 4C). A marked decrease in the survival of cMPL-expressing cells was observed after 48 hours of treatment with DT390-biscFV(cMPL), with an inhibitory concentration 50% (IC50) value of 21 pM, indicating a significant cMPL-mediated cytotoxic response compared to native cMPL-HEK293A cells (Figures 4D, E). Additionally, the effect of DT390-biscFV(cMPL) on the proliferation of human HSPCs was examined by segregating MPB CD34+ cells into cMPL^high^ and cMPL^low^ subsets using fluorescence-activated cell sorting. These subsets were then exposed to increasing concentrations of DT390-biscFV(cMPL) or the control immunotoxin (Figure 4C). After six days of culture, a significant decline in cell counts was observed, particularly within the cMPL^high^CD34+ group (IC50 = 98 nM), compared to the cMPL^low^CD34+ fraction (IC50 = 246 nM) (Figures 4F, G). These *in vitro* findings underscore the potential of DT390-biscFV(cMPL) to selectively target and eliminate human cMPL-expressing cells.

### DT390-biscFV(cMPL) depletes human cMPL+CD34+ cells in a xenograft mouse model

We next examined the *in vivo* efficacy of DT390-biscFV(cMPL) in depleting human HSPCs expressing cMPL. Utilizing a previously established protocol^15^, immunodeficient NBSGW mice were ‘humanized’ by transplantation of 1 x 10^6^ human MPB CD34+ cells. Following a 12-week period post-transplantation, which allowed for the establishment of stable hematopoiesis, the mice were administered either a single maximum tolerated dose of 1.2 mg/kg of DT390-biscFV(cMPL) or a PBS vehicle control (Supplemental Figure 3A). Serial PB sampling post-administration revealed no significant reduction in HSPC activity in the DT390-biscFV(cMPL)-treated mice (Supplemental Figures 3B, C). Additionally, at the conclusion of the study, no significant differences were observed in human cell engraftment within the BM and SP between the DT390-biscFV(cMPL) and PBS-treated groups (Supplemental Figures 3D-G). To address the possibility that the high levels of human cell engraftment observed following the transplantation of 1 x 10^6^ human MPB CD34+ cells could mask differences between the experimental groups, we modified our methodology by implementing a tenfold reduction in the number of transplanted human MPB CD34+ cells. Nevertheless, even with this revised protocol, the outcomes closely aligned with those of the original experiment, reinforcing the initial observations (data not shown).

To explore the underlying reasons for the lack of human HSPC depletion by DT390-biscFV(cMPL) in our xenograft model, we hypothesized that the expression of the human cMPL receptor might be suboptimal within the murine BM microenvironment. This hypothesis arose from the known variances in affinity and biological activity between murine and human cytokines, including TPO. To test this, we ‘humanized’ NBSGW mice with 1 x 10^6^ human MPB CD34+ cells and monitored cMPL expression on these engrafted human CD34+ cells within the murine BM over a 12-week period post-transplantation. Supporting our hypothesis, a progressive decrease in cMPL expression on these cells was observed (Supplemental Figures 3H-J).

To optimize the timing of DT390-biscFV(cMPL) administration, we treated ‘humanized’ NBSGW mice with the immunotoxin or a PBS control one day post-transplantation (Figure 5A). This timing was chosen based on the detectable surface expression of cMPL on human CD34+ cells newly homed to the murine BM. We then monitored human cell engraftment at early (1 week) and later (16 weeks) stages post-transplantation. At the one-week mark, a significant two-fold decrease in the percentage of human cMPL+CD34+ cells was observed in the BM of DT390-biscFV(cMPL)-treated mice compared to controls, suggesting effective targeting and depletion of the newly engrafted cells (Figure 5B). This finding was corroborated by a notable reduction in human cell engraftment in both the BM (Figures 5C, D) and SP (Figures 5E, F) of treated mice at 16 weeks. Collectively, these results not only underscore the specific targeting ability of DT390-biscFV(cMPL) towards human cMPL+CD34+ cells but also highlight its effectiveness in accessing the BM niche and depleting these cells in a live organism.

**Figure 5.**
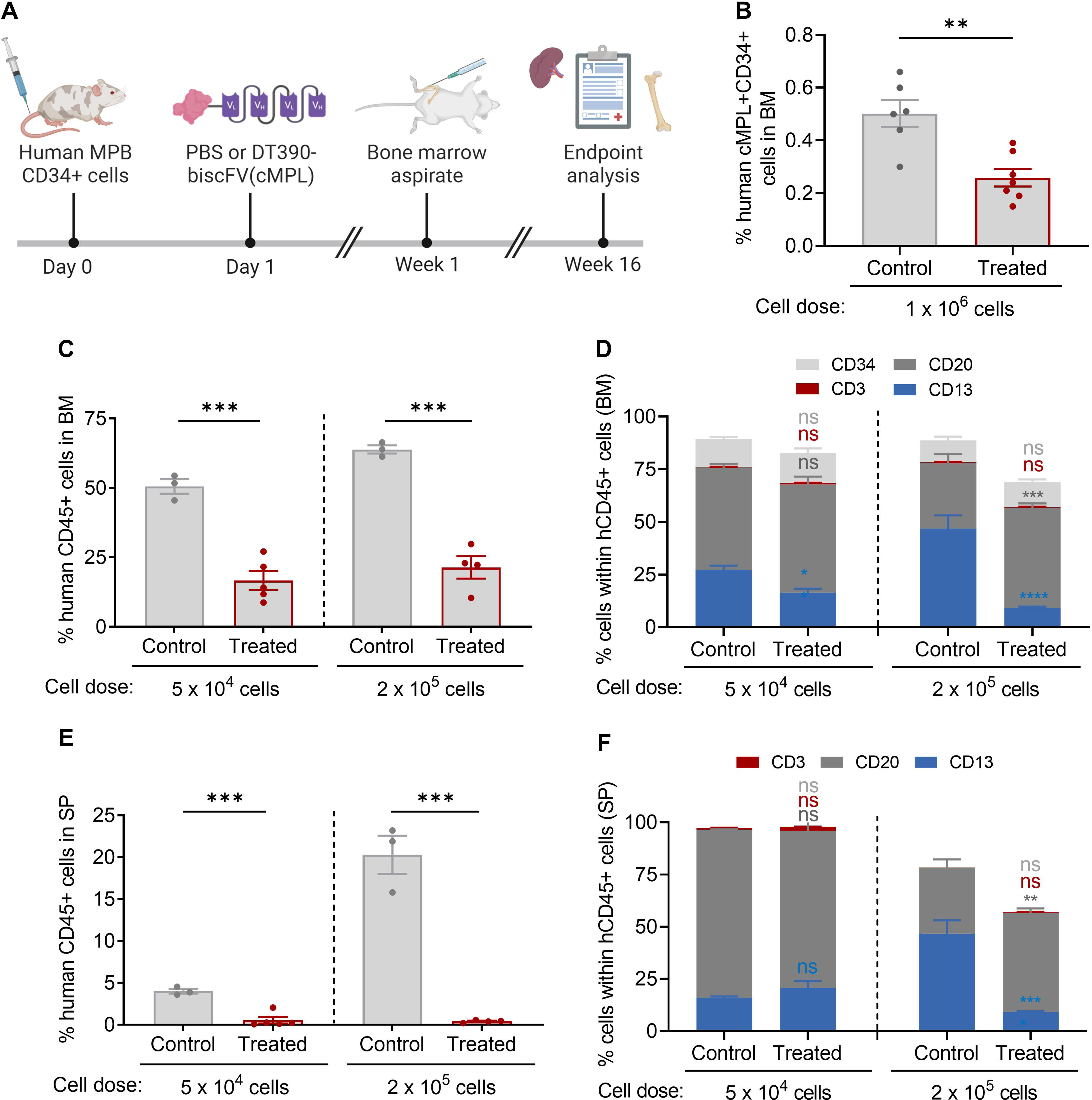
DT390-biscFV(cMPL) depletes human cMPL+ HSPCs in a xenograft mouse model. (A) Schematic of the experimental procedure: NBSGW mice were transplanted with human mobilized peripheral blood (MPB) CD34+ cells, followed by a single maximum tolerated dose of 1.2 mg/kg DT390-biscFv(cMPL) or PBS administered one day post-transplantation. Murine bone marrow (BM) was initially assessed at 1 week post-transplantation (cell dose: 1 x 10^6^), and subsequently both murine BM and spleen (SP) were evaluated at 16 weeks post-transplantation (cell dose: 5 x 10^4^ or 2 x 10^5^). (B) Percentage of human cMPL+CD34+ cells within the murine BM one week after administration of PBS (control) or DT390-biscFV(cMPL) (treated) (n = 5-6 mice per group). (C, D) Percentage of human CD45+ cells (C) and their lineage distribution (D) within the murine BM at the 16-week endpoint analysis for control and treated groups across both transplantation cell doses (n = 3-5 mice per group). (E, F) Percentage of human CD45+ cells (E) and their lineage distribution (F) within the murine SP at the 16-week endpoint analysis for control and treated groups across both transplantation cell doses (n = 3-5 mice per group). Datapoints represent the mean ± standard error of the mean (SEM), with statistical significance assessed via two-sided unpaired t-tests (panels B, C and E) or two-way ANOVA with Sidak multiple comparisons tests (panels D, F). Notations of statistical significance are as follows: ns, not significant, ** p ≤ 0.01, *** p ≤ 0.001, **** p ≤ 0.0001.

### DT390-biscFV(cMPL) enables safe and effective depletion of cMPL^high^CD34+ cells in rhesus macaques

To assess the translational potential of DT390-biscFv(cMPL), its efficacy and safety were investigated in rhesus macaques, a preclinical model with close phylogenetic ties to humans. This model recapitulates human *in vivo* TPO:cMPL signaling dynamics, providing a robust predictor for human gene therapy outcomes. Our initial step involved a cross-reactivity analysis of anti-human cMPL antibodies, leading to the identification of REA250 and 1.6.1 clones as the most efficient in recognizing rhesus cMPL receptors on CD34+ cells (Supplemental Figure 4A). Further studies confirmed a higher density of cMPL receptors on rhesus long-term NHPs LT-HSCs, identified as CD34+CD38-CD90+CD45RA-CD49f+, compared to the broader CD34+ HSPC population, paralleling human HSPC cMPL expression patterns (Figure 6A).

**Figure 6.**
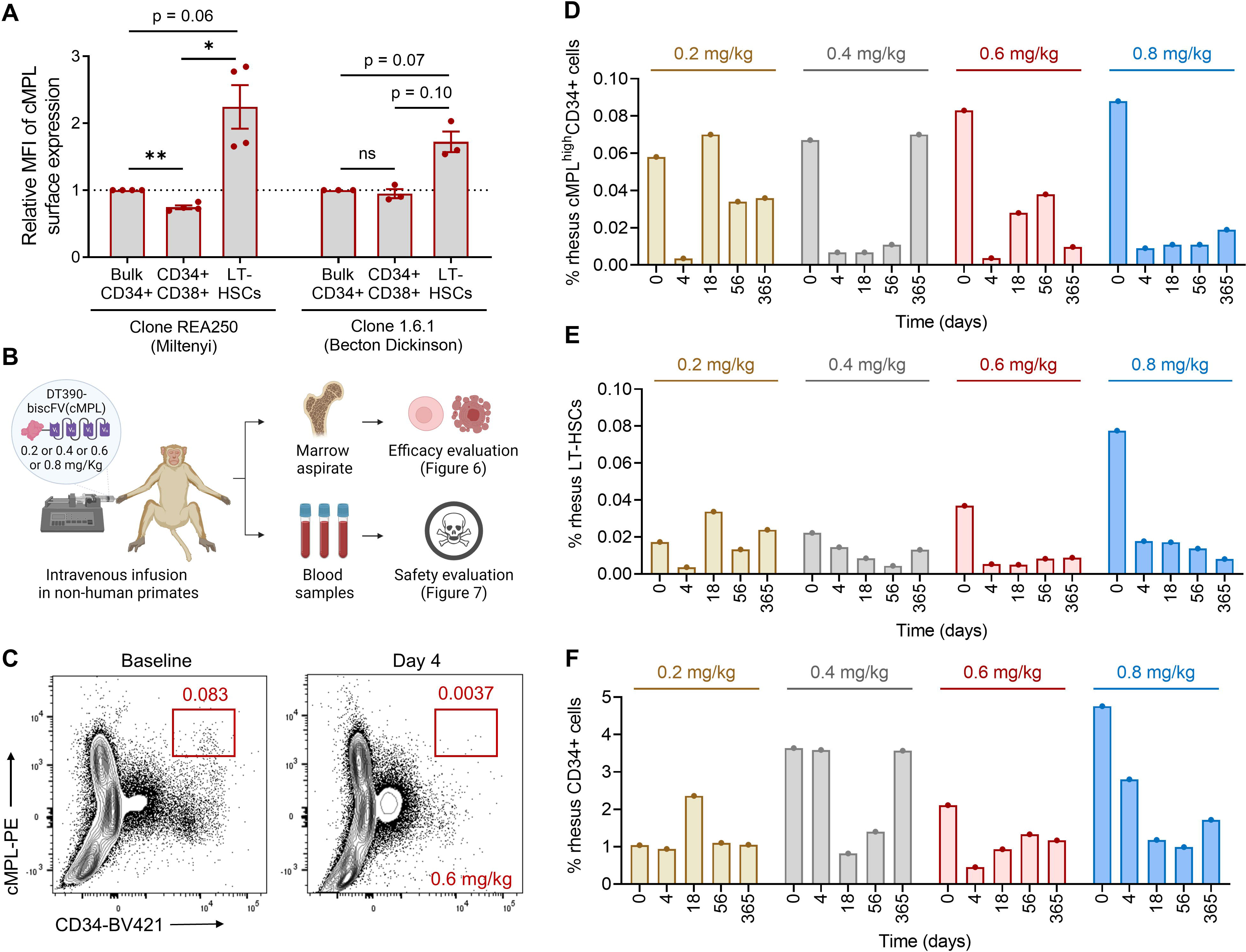
DT390-biscFV(cMPL) preferentially depletes LT-HSCs in rhesus macaques. (A) Relative mean fluorescence intensity (MFI) of cMPL expression on the cell surface within specified hematopoietic populations, normalized to the MFI of cMPL on bulk CD34+ cells (n = 4 animals per group). (B) Schematic of the experimental procedure: a single intravenous infusion of DT390-biscFv(cMPL) was administered at dosages of 0.2, 0.4, 0.6, or 0.8 mg/kg to four separate rhesus macaques. Post-administration, the impact on hematopoietic cell populations was monitored through serial sampling of bone marrow (BM) and peripheral blood. (C) Representative flow cytometry plots exemplifying the cMPL^high^CD34+ cell populations within BM aspirates pre-treatment (baseline) and four days after a 0.6 mg/kg dose of DT390-biscFv(cMPL). (D-F) Frequency of primitive hematopoietic subsets in BM samples collected pre-treatment and on days 4, 18, 56, and 365 following treatment (n = 1 animal per DT390-biscFV(cMPL) dose tested). Hematopoietic populations displayed include cMPL^high^CD34+ cells (D), LT-HSCs (characterized as CD34+CD38-CD90+CD45RA-CD49f+ cells) (E), and bulk CD34+ cells (F). Datapoints in panels A, and D-F represent the mean ± standard error of the mean (SEM). In panel (A), statistical significance was assessed via one-way ANOVA with Tukey correction. Notations of statistical significance are as follows: ns, not significant, * p ≤ 0.05, ** p ≤ 0.01.

*In vitro* experiments demonstrated the effective eradication of rhesus CD34+ HSPCs by DT390-biscFv(cMPL) (Supplemental Figures 4B, C). Subsequently, *in vivo* studies were conducted to evaluate the immunotoxin’s efficacy, safety, and pharmacokinetics in rhesus macaques. Following intravenous administration of single doses ranging from 0.2 to 0.8 mg/kg of DT390-biscFv(cMPL), HSPC levels were monitored through immunophenotyping of BM samples pre-treatment and on days 4, 18, 56, and 365 (Figure 6B). Within four days, a profound >90% reduction in the cMPL^high^CD34+ cell population was observed across all dosage levels (Figures 6C, D), with a more pronounced and lasting depletion of the phenotypically defined LT-HSC population (Figure 6E) compared to the entire CD34+ cell pool (Figure 6F). This observation suggests a dose-dependent mechanism of action, where lower doses yielded a temporary effect, whereas higher doses achieved a more sustained depletion of these cells, lasting up to one year (Figures 6D-F).

Pharmacokinetic analysis using an indirect ELISA revealed a notably brief half-life of DT390-biscFv(cMPL), ranging from 33 to 163 minutes and exhibiting dose-dependent kinetics (Figure 7A). This suggests a dose-responsive saturation of cMPL receptors, extending the immunotoxin’s presence in the circulation. The immunotoxin was undetectable by 24 hours, indicating rapid clearance, which is advantageous for subsequent cell reinfusion post-conditioning.

**Figure 7.**
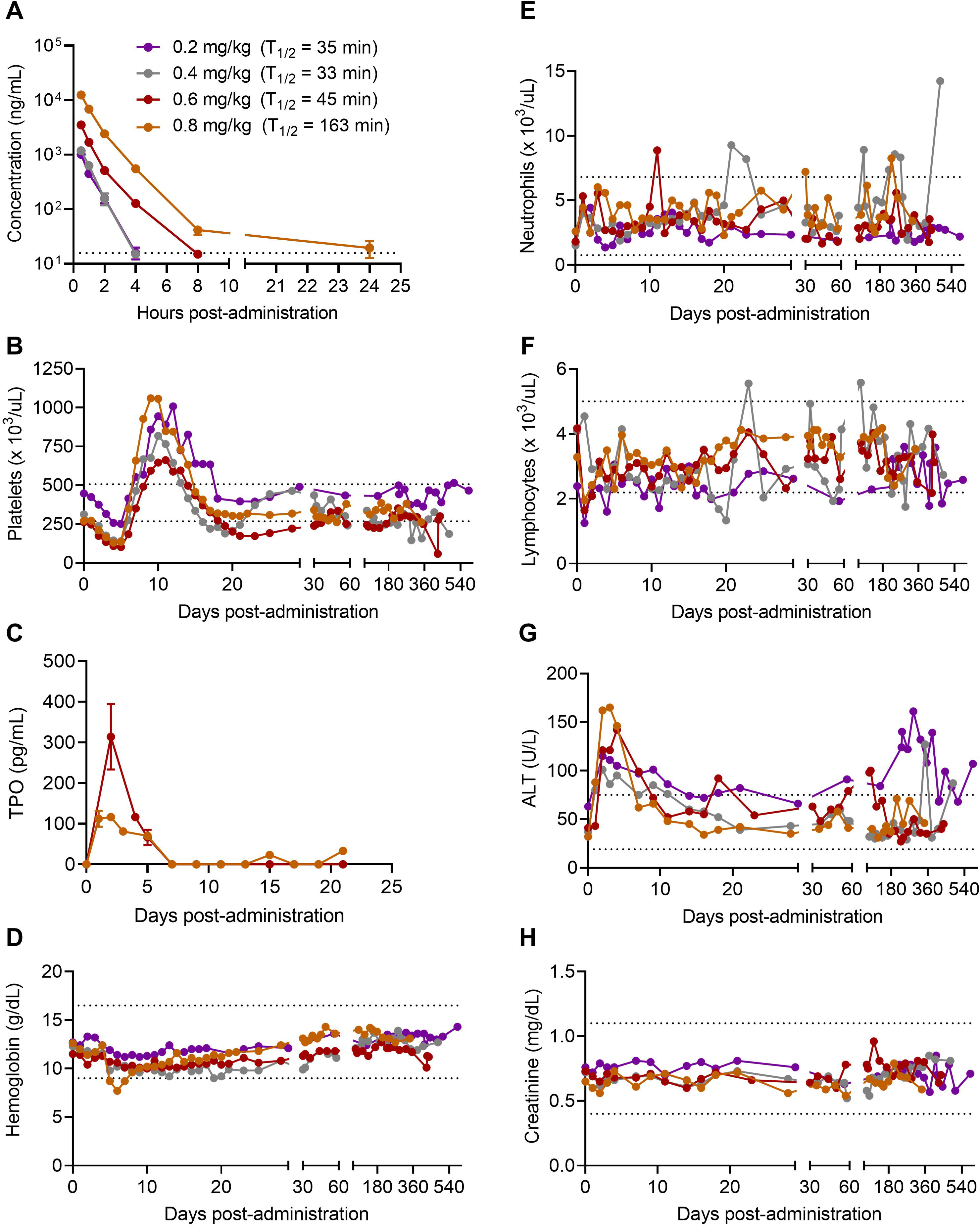
Safety and Pharmacokinetics Profile of DT390-biscFV(cMPL) in rhesus macaques. (A) Pharmacokinetic profile: The graph illustrates the time-dependent serum concentration of DT390-biscFV(cMPL) post-administration (n = 3 technical replicates at each timepoint). The dotted line represents the assay’s sensitivity threshold at 15 ng/mL. Note that the drug’s concentration falls below the detectable limit within 24 hours, highlighting its short systemic half-life. Legend in panel A applies to all panels. (B) Temporal changes in platelet counts following DT390-biscFV(cMPL) administration. (C) Luminex assay data indicating the serum thrombopoietin (TPO) levels post-treatment (n = 2 technical replicates at each timepoint).(D) Hemoglobin levels after treatment. (E) Neutrophil trajectory after treatment. (F) Lymphocyte counts after treatment. (G) Alanine transaminase (ALT) levels measured after treatment with DT390-biscFV(cMPL) to assess hepatic function and potential drug-induced hepatotoxicity. (H) Creatinine levels measured after treatment with DT390-biscFV(cMPL) to evaluate renal function and potential drug-induced nephrotoxicity. The dotted lines depicted in panels B and D-H represent the established normal range for each blood parameter measured. Datapoints in panels A and C represent the mean ± standard error of the mean (SEM).

Safety assessments included clinical monitoring and serial blood sampling. The animals maintained normal behavior and activity levels, with a transient, mild-to-moderate thrombocytopenia observed, reaching its nadir on days 4-5 and then showing robust recovery, exceeding baseline levels by days 9-12 (Figure 7B). This post-treatment thrombocytosis, attributed to a surge in endogenous TPO levels (Figure 7C), normalized rapidly as circulating TPO levels decreased due to sequestration by newly produced platelets and megakaryocytes. No adverse effects were noted on other hematopoietic lineages, with stable hemoglobin levels and neutrophil and lymphocyte counts (Figures 7D-F). A transient increase in liver alanine transaminase (ALT) levels was observed, but this was reversible and without lasting hepatic effects (Figure 7G). Electrolyte balance and renal function (Figure 7H) remained within normal ranges.

In summary, these *in vivo* findings in a rhesus macaque model demonstrate that DT390-biscFv(cMPL) is a targeted, dose-responsive agent that can safely and effectively deplete cMPL^high^ CD34+ cells. This underscores its potential clinical significance in conditions requiring the selective elimination of these cell populations.

## Discussion

The importance of the TPO:cMPL axis in early hematopoiesis, initially documented in murine HSPCs, has gained increasing recognition in human hematopoiesis. The development of small molecule agonists targeting the cMPL receptor, such as eltrombopag, has proven efficacious in maintaining and expanding human HSCs *ex vivo*^44,45^, as well as in restoring trilineage hematopoiesis in patients with immune BM failure syndromes^32–35^. Additionally, a comprehensive analysis of human hematopoietic lineage development demonstrated that incorporating cMPL (CD110) and CD71 into a conventional seven-surface marker phenotype elucidates cellular heterogeneity within early hematopoietic common myeloid progenitors (CMPs), megakaryocyte-erythroid progenitors (MEPs), and multipotent progenitors (MPPs), thereby challenging and redefining the traditional model of human HSC differentiation^46^.

However, the efficacy of cMPL as a singular marker for the identification of LT-HSCs within the adult human CD34+ cell population has not been demonstrated. Here, through extensive phenotypic, molecular, and functional analyses in xenograft models, we have established that high surface expression of cMPL alone is a robust marker for the enrichment of LT-HSCs within the human CD34+ cell pool. Employing a conservative sorting approach to segregate cMPL^high^ cells (top 10%) from cMPL^low^ cells (bottom 10%), we observed a frequency of 1 LT-HSC per 1,266 CD34+ cells, indicating a substantial 152-fold enrichment of LT-HSCs within the cMPL^high^CD34+ cell subset relative to cMPL^low^CD34+ cells. Refining the sorting strategy to selectively target CD34+ cells that express cMPL at levels surpassing those utilized in our study could further enhance the enrichment of LT-HSCs, potentially achieving near single-cell precision. The cMPL^high^CD34+ subset distinguishes LT-HSCs from their more differentiated progeny, offering a streamlined combination of positive markers for biological, molecular, and therapeutic investigations focused on the human HSC population. Our studies in the NHP large animal model reinforce the specificity and significance of cMPL receptor expression within the LT-HSC population, as evidenced by the preferential depletion of LT-HSCs over the total CD34+ cell population following the administration of a cMPL-specific immunotoxin.

The DT390-biscFv(cMPL) immunotoxin has several key attributes enhancing its specificity and effectiveness in LT-HSC depletion, while also ensuring increased safety *in vivo*. First, the agonistic properties of the biscFV(cMPL) moiety enhance LT-HSC depletion by facilitating cMPL dimerization and internalization upon binding^40^. Moreover, the absence of antibody constant fragments (Fc) reduces non-specific Fc-mediated cellular binding, thereby minimizing off-target effects. This construct has also been modified to exclude the native DT binding domain, preventing internalization in non-target cells. Importantly, this truncated form of DT circumvents inhibition by pre-existing antibodies in human blood^47^. Considering the immunogenicity concerns in therapies necessitating multiple administrations, the inclusion of biscFv(cMPL), a humanized minibody, substantially reduces this risk if repeated applications of the immunotoxin are required^40^. The FDA approval of other DT390-based fusion toxins for repeated treatments, such as Ontak^48^ and Elzonris^49^ targeting IL-2 and IL-3, respectively, suggests a favorable outlook for the clinical tolerance of DT390-biscFv(cMPL) in therapeutic settings. Lastly, DT390-biscFV(cMPL) displays a remarkably shorter half-life compared to monoclonal antibodies targeting cKIT, a feature potentially advantageous in clinical scenarios necessitating rapid immunotoxin clearance^20^.

DT390-biscFv(cMPL)’s high specificity in LT-HSC depletion may unveil novel scientific insights into early hematopoiesis and broaden the therapeutic scope of TPO:cMPL manipulation beyond the conventional focus on activating this pathway for HSC maintenance and expansion. DT390-biscFV(cMPL) emerges as a promising candidate for pre-transplant conditioning, primarily in *ex vivo* autologous gene therapy for patients with inherited, non-malignant disorders, where preservation of immunity is paramount. In particular, it may be well-suited in congenital BM failure syndromes like Fanconi anemia, by targeting and eliminating residual host LT-HSCs prone to malignant transformation. Additionally, DT390-biscFV(cMPL)-based conditioning regimens may offer valuable insights when comparing transplantation outcomes between enriched LT-HSCs and bulk CD34+ cell populations. Enriching LT-HSCs from human CD34+ HSPCs can lower costs and improve gene editing efficiency in *ex vivo* gene therapies^50–56^, but this process also significantly reduces hematopoietic progenitors, potentially delaying early hematopoietic reconstitution and necessitating frequent transfusions post-transplant due to cytopenias^53^. DT390-biscFV(cMPL) could address this by selectively depleting LT-HSCs while preserving progenitors, thus enabling safer exploration of LT-HSC purification in gene therapy contexts. In allogeneic HSPC transplantation, combining DT390-biscFv(cMPL) with lymphocyte depleting drugs to prevent immune-mediated rejection of the donor graft may also offer significant advantages over traditional conditioning methods.

The findings from our study highlight multiple avenues for future research. Previous investigations targeting the cKIT receptor in host HSPCs underscore the feasibility of precise HSPC eradication as a conditioning strategy for transplantation^13,15^. However, further investigations are needed to validate whether DT390-biscFV(cMPL)’s enhanced specificity for LT-HSCs, as compared to anti-cKIT strategies, provides adequate niche availability for donor HSC engraftment. Future work will investigate the effects of increasing DT390-biscFV(cMPL) doses on LT-HSC populations and assess hematopoietic and non-hematopoietic toxicities. These studies will also aim to identify the optimal DT390-biscFV(cMPL) administration protocol (e.g., single dosing vs. split dosing), the most suitable HSPC transplantation timeframe, and the synergistic use of DT390-biscFV(cMPL) with other chemotherapy-free conditioning methods. Notably, the potential of using mobilizing agents as a conditioning regimen^57,58^ offers a promising avenue for integration with DT390-biscFV(cMPL) to selectively eliminate mobilized LT-HSCs and enhance the engraftment of transplanted cells. Our work will also facilitate further exploration into the diversity and hierarchical organization within the phenotypically distinct HSC compartment^59,60^. Specifically, a subset of primitive TPO-dependent HSCs predisposed to platelet production has been identified in murine models^39,61–64^. These cells are positioned at the apex of the HSC hierarchy and have demonstrated capacities for both transient and stable long-term platelet-biased multilineage reconstitution. The prospect that DT390-biscFV(cMPL) may specifically target this distinct cell population warrant further investigations.

In conclusion, the current study not only elucidates the critical role of the TPO:cMPL axis in HSC biology but also introduces DT390-biscFv(cMPL) as a novel and potent tool for specific LT-HSC depletion. Our findings highlight the potential of DT390-biscFv(cMPL) in conditioning regimens for both autologous and allogeneic transplants. Future research will focus on optimizing DT390-biscFv(cMPL) dosage and administration protocols, assessing its synergistic potential with other conditioning strategies, and further investigating its targeted effects on specific HSC subpopulations, thereby advancing our understanding and therapeutic manipulation of human hematopoiesis.

## STAR ⍰ METHODS

### KEY RESOURCES TABLE

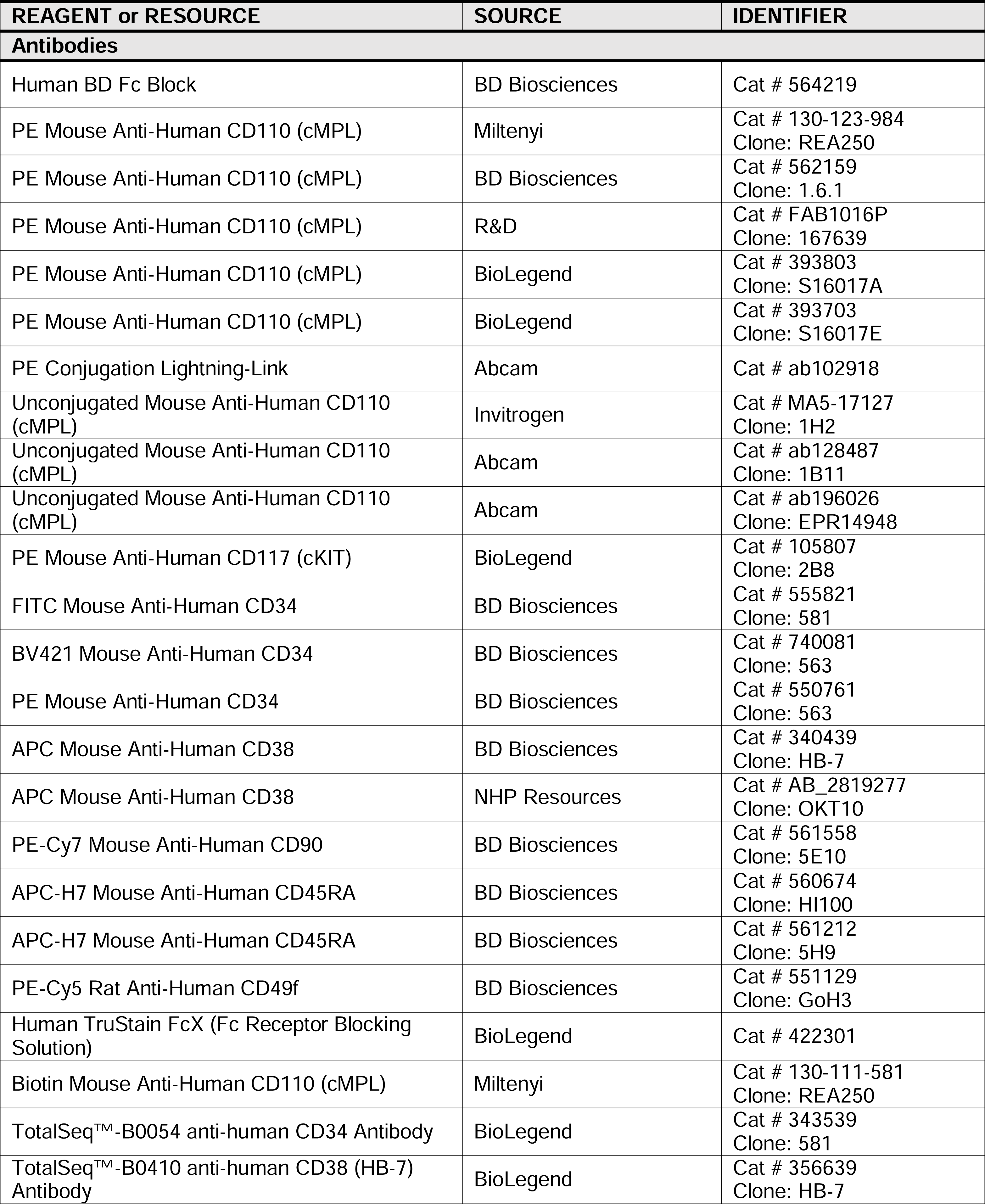
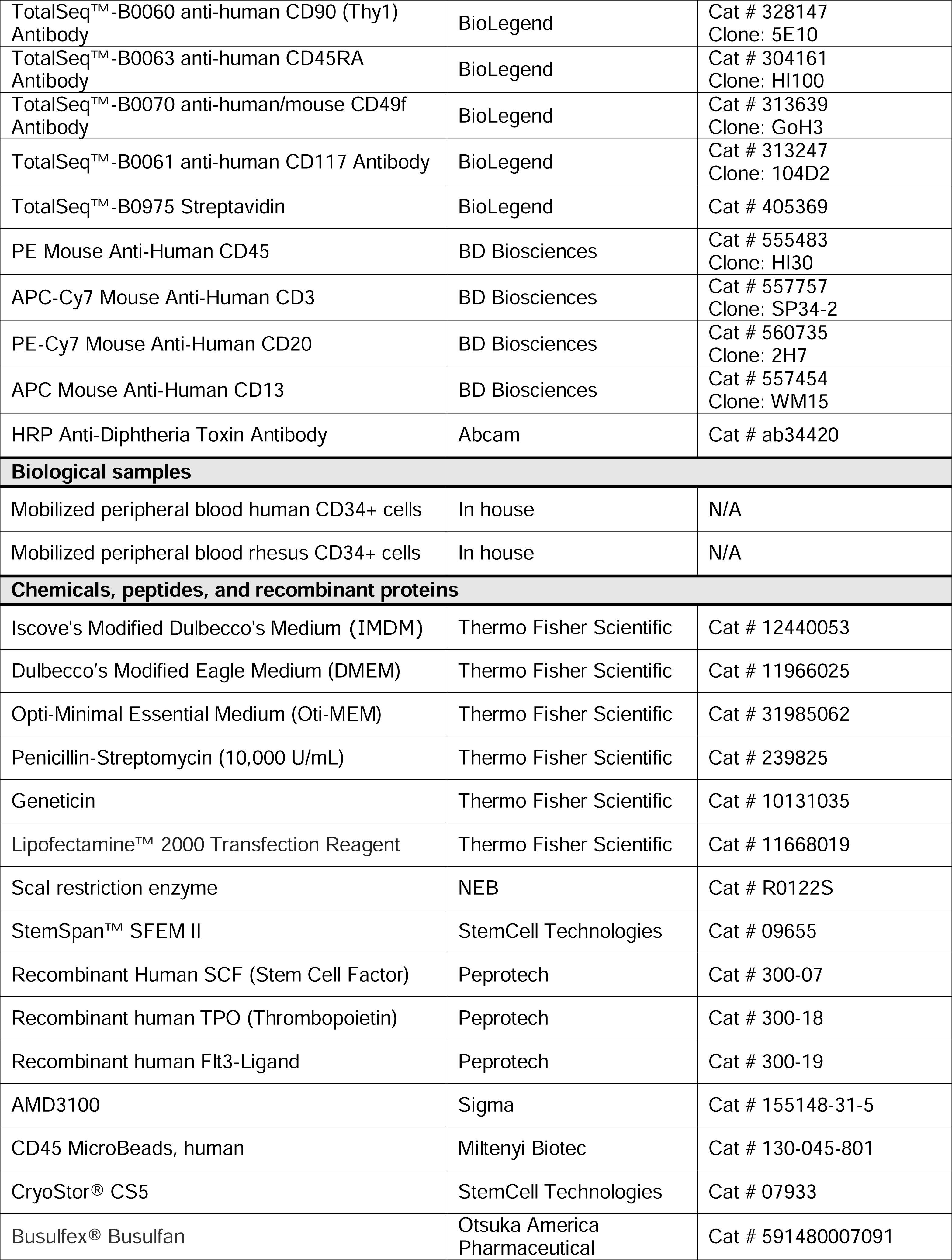
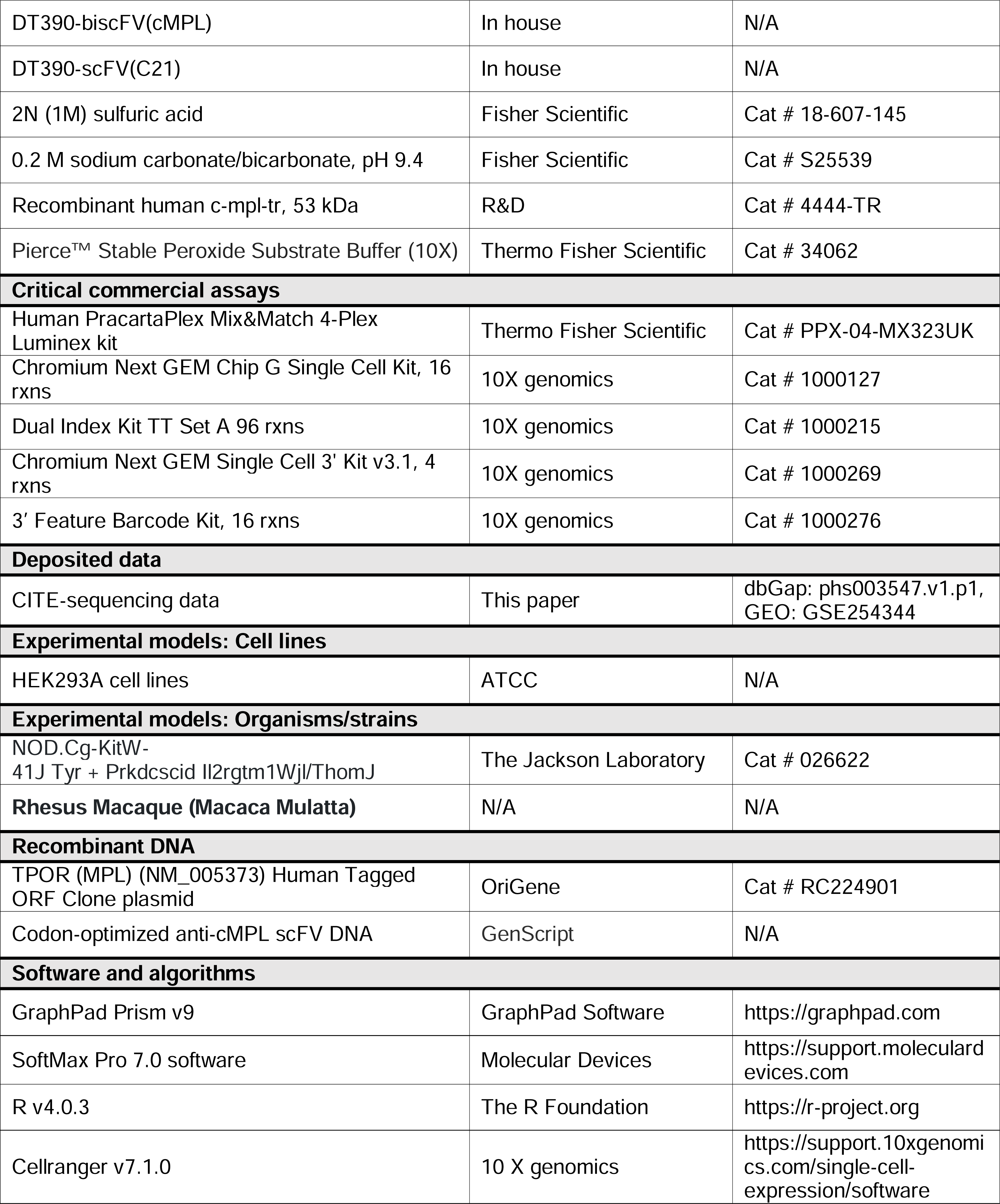

## RESOURCE AVAILABILITY

### Lead contact

Further information and requests for resources should be directed to Andre Larochelle, MD., Ph.D.

### Materials availability

This study generated a truncated diphtheria toxin-based recombinant anti-cMPL immunotoxin (i.e., DT390-biscFV(cMPL)). The patent number is E-188-2021-0.

### Data and code availability

CITE-sequencing data are available at dbGap via authorized access request as detailed at https://dbgap.ncbi.nlm.nih.gov/aa/wga.cgi?page=login (dbGap accession number: phs003547.v1.p1). Data are also available in Gene Expression Omnibus under accession GSE254344.

## EXPERIMENTAL MODELS AND SUBJECT DETAILS

### Studies involving humans and animals

For human studies, CD34+ HSPCs were procured from healthy volunteers under informed consent in strict adherence to the Declaration of Helsinki. This process was conducted under a clinical protocol sanctioned by the Institutional Review Board (IRB), bearing the identifier NCT00001529. For murine and rhesus macaque studies, all procedures complied with the guidelines set by the Committee on Care and Use of Laboratory Animals of the Institute of Laboratory Animal Resources, National Research Council (DHHS publication No. NIH 85-23), and were approved by the Animal Care and Use Committee of the National Heart, Lung, and Blood Institute.

### Isolation of human CD34+ HSPCs

Human HSPCs were mobilized in the PB of healthy donors, aged 18 to 60 and spanning both genders, through a five-day regimen of subcutaneous Granulocyte-Colony Stimulating Factor (G-CSF) (Filgrastim, Amgen, Thousand Oaks, CA, USA) injections at a dose of 10 μg/kg. This was followed by leukapheresis, conducted using the Cobe Spectra Apheresis System (Terumo BCT, Lakewood, CO, USA). The leukapheresis product, primarily mononuclear cell concentrates, underwent enrichment for CD34+ HSPCs utilizing the CliniMACS Plus instrument (Miltenyi Biotec, Gaithersburg, MD, USA), and were subsequently cryopreserved for future use.

### Isolation of rhesus CD34+ HSPCs

The isolation of rhesus CD34+ HSPCs was conducted on MPB, obtained following a cytokine mobilization regimen comprising a five-day course of 15 - 20 mcg/kg/day G-CSF (Amgen) and a single dose of 1 mg/kg AMD3100 (Sigma-Aldrich, MO, USA) administered subcutaneously^69^. Leukapheresis, using a CS3000 Cell Separator (Baxter Fenwal, IL, USA), was performed 2 to 4 hours after the AMD3100 administration. Rhesus PBMCs were collected using Ficoll-Paque PLUS density gradient media (GE Healthcare, Sweden) following the manufacturer’s instructions. Immunoselection of CD34+ HSPCs from the leukapheresis product was accomplished using a rhesus-specific CD34+ antibody (Clone 12.8, Fred Hutchinson Cancer Research Center, USA) and anti-mouse IgM beads (Miltenyi Biotech). The purity of the selected cells was confirmed using an additional CD34+ antibody (Clone 563, BD) and 7-AAD staining. These cells were then cryopreserved for subsequent experiments.

## METHOD DETAILS

### Analysis of cMPL and cKIT Surface Expression in CD34+ HSPCs

Freshly thawed human and rhesus CD34+ cells were incubated with human Fc Block^TM^ (Becton Dickinson (BD), Franklin Lakes, NJ, USA, 564219) for 10 minutes at room temperature (RT) to minimize non-specific antibody interactions. These cells were then subjected to a 30-minute staining procedure on ice (all antibodies used in 1:25 dilution), utilizing two distinct antibody panels with or without PE-anti-cMPL (Miltenyi Biotech) or PE-anti-cKIT (BioLegend, San Diego, CA, USA, 105807). The human HSC panel included FITC-anti-CD34 (BD, 560942), APC-anti-CD38 (BD, 340439), PE-Cy7-anti-CD90 (BD, 561558), APC-H7-CD45RA (BD, 560674), and PE-Cy5-anti-CD49f (BD, 551129) antibodies. The rhesus HSC panel comprised BV421-anti-CD34 (BD, 740081), APC-anti-CD38 (NHP Resources), PE-Cy7-anti-CD90 (BD, 561558), APC-H7-CD45RA (BD, 561212) and PE-Cy5-anti-CD49f (BD, 551129) antibodies. In a specified set of experiments, antibodies underwent conjugation employing a PE conjugation lightning-link kit (Abcam, Waltham, MA, USA, ab102918). A detailed inventory of the antibodies utilized is available in Key Resources Table. Stained cells were subjected to a single wash with FACS buffer and then filtered through 40 µm cell strainers. Quantitative and qualitative analyses of surface marker expression were then conducted using an LSR II Fortessa flow cytometer (BD).

### CITE-sequencing

The 3’ end scRNA-seq was performed on a Chromium Single-Cell Controller (10X genomics, Pleasanton, CA, USA) using the chromium Single Cell 3’ Reagent Kit v3.1 dual index kit according to the manufacturer’s instructions. Human MPB CD34+ cells from a healthy volunteer were thawed using chilled Iscove’s Modified Dulbecco’s Medium (IMDM) (Thermo Fisher Scientific, Waltham, MA, USA,) supplemented with 2% fetal bovine serum (FBS), washed twice, and counted with AOPI staining on an automated cell counter. A total of 1 x 10^6^ cells were resuspended in 10 uL of PBS supplemented with 2% FBS and 2 mM EDTA (hereafter named ‘FACS buffer’) together with 5 uL of TruStain FcX (BioLegend, 422301). Cells were incubated with the FcX block for 10 minutes at RT, after which cells were stained with biotin-conjugated-anti-cMPL antibody (Miltenyi Biotech 130-111-660, used in 1:25 dilution) on ice for 30 minutes. Cells were washed twice and blocked again with TruStain FcX. Subsequently, 0.5 ug each of the 10 TotalSeq-B oligo-tagged antibodies (BioLegend) listed in Key Resources Table were added and cell pellets were suspended. After incubation with TotalSeq antibodies on ice for 30 minutes, cells were washed three times and approximately 2.2 x 10^4^ cells were utilized for the subsequent procedure.

Oligo-tagged cells were loaded on a Chromium Chip B with master mix, gel beads, and partitioning oil, again according to 10x Genomics instructions. 3’ v3.1 GEX libraries were constructed according to 10x Genomics protocol. 3’ v3.1 GEX and ADT libraries were pooled in a 2:1 ratio and sequenced with an Illumina NovaSeq S1 flow cell (100 cycles, 2 lanes). Target sequencing depth for the GEX libraries was 2 x 10^4^ read pairs per cell and 5 x 10^3^ read pairs per cell for the ADT libraries.

Illumina sequencer’s base call files (BCLs) were demultiplexed, for each flow cell directory, into FASTQ files using Cellranger ‘mkfastq’ with default parameters (v 7.1.0) except ‘include-introns’ option set to ‘false’ on NIH Helix/Biowulf High Performance Computing Cluster (HPC). FASTQ files were then processed using Cellranger ‘count’ with default parameters. Internally, the software relies on STAR for aligning reads to a pre-build GRCh38 human reference genome, while genes are quantified using ENSEMBL genes as gene model. The output of Cellranger is a filtered gene-barcode matrix containing the UMI counts for each gene.

Gene counts were processed with Seurat (v 4.1.0, https://satijalab.org/seurat/). Quality control was performed to filter low-quality cells by excluding cells expressing genes more than [mean + 2 x standard deviations (SD)] of total genes, less than [mean - 2 x SD] of total genes, or mitochondrial genes comprising more than 10% of the total gene expression per cell. Counts were normalized using Seurat function NormalizeData with default parameters. Expression data were then scaled using the ‘ScaleData’ function. The doublets were identified using scDblFinder (v 1.4.0) for further filtering. To decrease noise and make downstream computations more tractable, the dataset was reduced to 20 dimensions of PCA using the ‘RunPCA] function. For visualization, the dataset was dimensionally reduced to two dimensions of Uniform Manifold Approximation and Projection (UMAP) using ‘RunUMAP function. Shared-nearest neighbor (SNN) graph based on cell-cell distance matrix constructed using ‘FindNeighbors’ function was subsequently used to partition cells into clusters using Louvain modularity optimization algorithm using FindClusters [function at resolution of 0.3. This resulted in 7 distinct HSC and progenitor clusters.

We first evaluated expression levels of lineage defining genes within these clusters: highest expression of EGR1, MLLT3 and HES1 genes in the HSC cluster 2; second highest expression of EGR1, MLLT3 and HES1 genes in the MPP cluster 3; GATA1, GATA2 and HBB expression in the MEP cluster 5; Myeloperoxidase (MPO), IFITM2 and IFITM3 expression in the LMPP cluster 0; IFITM2 and IFITM3 expression in the ETP cluster 4; DNTT and CD79A expression in the CLP cluster 6. CMP cluster 1 was annotated on the basis of being positioned between HSC/MPP/LMPP clusters and MEP cluster. Protein surface protein expression from CITE-seq data further confirmed that the transcriptionally defined HSC cluster 2 is consistent with the known HSC immunophenotype in humans, as evidenced by higher expression of CD90 and CD49f and lower expression of CD38 and CD45RA.

### Transplantation of CD34+ HSPCs into NBSGW mice

In the primary transplantation study, a specified quantity of CD34+ HSPCs was administered intravenously via tail vein injection into female NOD,B6.SCID IL-2r ^-^/^-^Kit^W41/W41^ (NBSGW) mice, aged between eight to sixteen weeks (Jackson Laboratory, Bar Harbor, ME, USA). Peripheral blood was sampled at predetermined intervals post-transplantation. For endpoint analysis at 16 weeks post-transplant, cells from PB and SP were evaluated using a specific antibody panel (PE-anti-CD45, PE-Cy7-anti-CD20, APC-anti-CD13 and APC-Cy7-anti-CD3 (all in 1:25 dilution)), and BM cells were assessed using a different antibody panel (PE-anti-CD45, PE-Cy7-anti-CD20, APC-anti-CD13, APC-Cy7-anti-CD3 and FITC-anti-CD34 (all in 1:25 dilution)). To isolate human CD45+ cells from the primary transplanted mice, the remaining fresh murine BM cells were purified using human CD45 MicroBeads (Miltenyi Biotech) and cryopreserved in CryoStor® CS5 (STEMCELL Technologies, Cambridge, MA, USA). For secondary transplantation, a 10 mg/kg dose of busulfan (Busulfex, Otsuka, Rockville, MD, USA) was administered intraperitoneally into female NBSGW mice 48 hours prior to tail-vein injection. This procedure was chosen based on previous studies indicating enhanced sensitivity for detecting human cell engraftment in this model^65^. Endpoint analysis in secondary transplants was performed at 16 weeks post-transplant. Bone marrow was harvested and assessed with an antibody panel that included PE-anti-CD45, PE-Cy7-anti-CD20, APC-anti-CD13 (all in 1:25 dilution). In the assessment of secondary mouse engraftment, a positive score was assigned when both myeloid (CD13+) and lymphoid (CD20+) lineages were clearly evident, coupled with the presence of human CD45+ cells exceeding 0.01% of the total cell population. The frequency of LT-HSCs was determined using a Limiting Dilution Analysis (LDA), as previously reported^66^. The HSC frequency was calculated using ELDA software (http://bioinf.wehi.edu.au/software/elda/) and plotted using the limdil function in the satmod package within RStudio IDE.

### Immunotoxin production

Bivalent anti-cMPL single-chain fragment variable (biscFV(cMPL)) (clone hVB22B) was previously described as a minibody demonstrating similar agonistic activity for human and NHP cMPLs^40^. Codon-optimized anti-cMPL scFV DNA was synthesized by GenScript (Piscataway, NJ, USA). DT390-biscFV(cMPL) was constructed, expressed and purified using a unique DT resistant yeast *Pichia pastoris* expression system^67^, as previously described^68^. DT390-scFV(C21), a DT390-based unrelated immunotoxin used as negative control, was also expressed and purified using the same DT resistant yeast *Pichia pastoris* expression system.

### *In vitro* culture of HEK293-A cells

The generation of human cMPL receptor-expressing HEK293-A cells was achieved through a lipofectamine-mediated transfection of a plasmid expressing cMPL and GFP (Origene), previously linearized using the Scal restriction enzyme (NEB). The cells, ensuring 70-90% confluence, were exposed to DNA-lipid complexes formed by a 1:1 mixture of Lipofectamine® 2000 Reagent (Thermo Fisher Scientific) and the aforementioned plasmid in Opti-MEM medium (Thermo Fisher Scientific). This mixture was allowed to interact at RT for 5 minutes prior to application. Post-transfection, the cells were cultured under 5% CO2 at 37°C for 48 hours. Subsequent selection involved serial dilutions in Opti-MEM medium containing Geneticin (Thermo Fisher Scientific, 10131035), ranging from 200-400 ug/mL, to isolate single clones positive for both cMPL and GFP, which were then expanded and cryopreserved for future use. For *in vitro* experimentation, 1 x 10^4^ cMPL receptor-positive and negative HEK293-A cells were seeded into 96-well plates, precoated for tissue culture, with each well containing 100 uL of Dulbecco’s Modified Eagle Medium (DMEM) (Thermo Fisher Scientific), supplemented with 1% penicillin-streptomycin (GIBCO, Grand Island, NY, USA) and 10% FBS. The cells were treated with varying concentrations of DT390-biscFV(cMPL) or control immunotoxins, maintained in an environment of 5% CO2 at 37°C. Viable cell enumeration was performed two days post-treatment using a Celigo image cytometer (Nexcelom Bioscience).

### *In vitro* culture of human CD34+ HSPCs

The *in vitro* culture of human CD34+ HSPCs involved plating 1 x 10^3^ G-CSF mobilized CD34+ cells or fluorescence-activated cell sorted cMPL^high^CD34+ cells in fibronectin-coated 96-well plates. Each well contained 100 uL of StemSpan SFEM II medium (STEMCELL Technologies) supplemented with 1% penicillin-streptomycin (GIBCO), 100 ng/mL SCF, and 100 ng/mL Fms-like tyrosine kinase 3 ligand (Flt3-ligand). Similar to the HEK293-A cell protocol, these cells were also treated with varied concentrations of DT390-biscFV(cMPL) or control immunotoxins, incubated at 5% CO2 and 37°C, with viable cell counts assessed after 6 days using a Celigo image cytometer.

### *In vitro* culture of rhesus CD34+ HSPCs

*In vitro* culture of rhesus CD34+ HSPCs involved plating 1 x 10^3^ G-CSF/AMD3100 mobilized cells in fibronectin-coated 96-well plates, each containing 100 uL of StemSpan SFEM II medium (STEMCELL Technologies), supplemented with 1% penicillin-streptomycin (GIBCO), 100 ng/mL SCF, 100 ng/mL Flt3-ligand, and 100 ng/mL TPO. The cells were treated with various concentrations of DT390-biscFV(cMPL) or control immunotoxins, incubated at 5% CO2 and 37°C, with cell viability assessed after 6 days using a Celigo cytometer.

### Administration of DT390-biscFV(cMPL) into rhesus macaques

On the day of treatment, Benadryl (2 mg/kg) was administered intramuscularly 30 minutes before the infusion of DT390-biscFV(cMPL). The animals received a 100 mL saline solution containing varying doses (0.2, 0.4, 0.6, or 0.8 mg/kg) of DT390-biscFV(cMPL), infused intravenously over 60 minutes. Clinical monitoring of the animals and their vital signs was conducted during and after the infusion.

### Collection of BM aspirates from rhesus macaques

BM aspirates from rhesus macaques were collected at multiple time points (days 0, 4, 18, 56, and 365 post-treatment). Under anesthesia, a small incision was made on the skin over the posterior iliac crests or ischial tuberosities, followed by the insertion of an 11-18-gauge BM needle into the marrow cavity. Bone marrow, up to 3 ml, was collected using a 10-mL heparin-coated syringe and resuspended in ∼10 ml PBS (pH 7.2 to 7.4) and heparin 3 (40 unit/ml) at RT, then filtered (70 µm) and processed for cell enumeration and mononuclear cell isolation.

### Indirect enzyme linked immunosorbent assay (ELISA)

After animals were anesthetized, plasma samples were collected at specific time points: 5 minutes before, and 30 minutes, 1, 2, 4, 8, 24, and 48 hours after DT390-biscFV(cMPL) administration, and stored at −80°C for later analysis. The immunoassay employed 96-well plates (ThermoFisher Scientific, Cat # 3455) coated with a 0.2 M sodium carbonate/bicarbonate buffer containing 5 µg/mL of recombinant human cMPL protein (R&D, Cat# 444-TR-050) and incubated overnight at 4°C. Following overnight incubation, plates were washed thrice for 5 minutes with 200 µL 1X PBS-T (ThermoFisher Scientific, Cat# 28352), and then blocked overnight at 4°C with 200 µL of 2% BSA in PBS-T to prevent non-specific binding. Following the blocking step, plates underwent an additional wash cycle, after which 100 μL aliquots of plasma samples, systematically prepared in a series of PBS-T dilutions at ratios of 1:2, 1:4, 1:8, 1:16, 1:32, 1:64, 1:128, and 1:248, were dispensed into the wells in triplicate. For standardization, wells containing various concentrations of DT390-biscFV(cMPL) in PBS-T (2000 ng/mL to 15.625 ng/mL) were also prepared. Plates were incubated overnight at 4°C. After another washing cycle, wells were incubated with 100 µL of 1 µg/mL anti-DT antibody conjugated to HRP (Abcam, Cat# ab34420) (1:1000 dilution) at RT for 1 hour. Following a final washing, a chromogenic reaction was initiated by adding 100 µL of stable peroxide buffer containing o-phenylenediamine dihydrochloride (OPD) substrate to each well, incubated for 30 minutes at RT. The reaction was halted using 50 µL of 2N (1M) sulfuric acid, and absorbance was measured at 492 nm using a SpectraMax Plus 384 Microplate Reader (Avantor) and SoftMax Pro 7.0 software. Plasma concentrations of DT390-biscFV(cMPL) at different timepoints were determined using a standard curve, analyzed with GraphPad Prism software.

### Luminex assay

Plasma TPO levels were estimated in plasma samples using Human PracartaPlex Mix&Match 4-Plex Luminex kit (ThermoFisher Scientific, Cat # PPX-04-MX323UK). Assays were performed as per manufacturer’s instructions. Briefly, samples were run in duplicates by incubating with cocktail of magnetic beads coupled with primary capture antibody for 2 hours at RT in dark on a plate shaker (825rpm). After incubation beads were washed 3 times and were incubated with biotinylated detection antibody cocktail for 1 hour at RT in dark on a plate shaker (825 rpm). Upon completion of incubations, wells were washed 3 times, before incubation with streptavidin PE solution in assay buffer for 30 mins at RT in dark on a plate shaker (825 rpm). Upon completion of incubation wells were washed 3 times and magnetic bead cocktail were resuspended in 125 uL of wash buffer provided in kit. Serial dilutions of standards were prepared as per kit’s recommendation to generate standard curve in pg/mL for respective analytes. concentrations of analytes in the test samples were extrapolated using Bioplex-manager 6.1 software on a Bioplex-200 Luminex reader (Bio-Rad).

### Statistics

Statistical analyses were conducted using GraphPad Prism 9 software, with data represented as mean ± standard error of the mean (SEM). Significance was determined using two-sided unpaired Student’s t-test, one-way ANOVA test with Tukey correction, or two-way ANOVA test with Sidak correction, with a significance threshold set at p < 0.05.

## Acknowledgements

The authors thank all members of the Larochelle Lab for invaluable discussions and assistance with murine BM harvesting; David Stroncek MD and the team at the NIH Department of Transfusion Medicine and Cell Processing Section for their expertise in apheresis, selection, and cryopreservation of human CD34+ cells; Richard Gustafson RN and the outpatient clinic nursing staff for their role in recruiting volunteers and instructing them on G-CSF administration; Aylin Bonifacino for NHP cell processing; the NIH Division of Veterinary Resources team as well as Building 49 Central Animal Facility staff for provision of excellent care for NHPs used in this study; Julie Holdridge DVM and the staff of Building 50 Animal Facility for their excellent care of the animals; Yuesheng Li, Yan Luo, Poching Liu and the NHLBI DNA Sequencing and Genomics Core staff for assistance with CITE-seq. This research received fundings from the Intramural Research Program of the NHLBI, NIH, USA (ZIA HL006172 and Z99 HL999999) and the NIH Office of the Director (P40 OD028116).

## Author contributions

D.A. and A.L. conceptualized the project and designed the experiments. D.A. performed experiments and acquired data. S.H., N.L., A.K., T.E. and J.G. performed NHP procedures and sample collections. D.A., B.F., and N.R. analyzed CITE-seq dataset. D.A. and C.S.R. harvested and processed murine BM and SP samples for flow cytometry analysis. D.A. and P.D. prepared samples for Luminex assay. D.A, D.M., Z.W. and A.L designed the construct of DT390-biscFV(cMPL). Z.W. produced and provided DT390-biscFV(cMPL). D.A. and A.L. wrote the original draft. All authors have read and agreed to the published version of the manuscript.

---

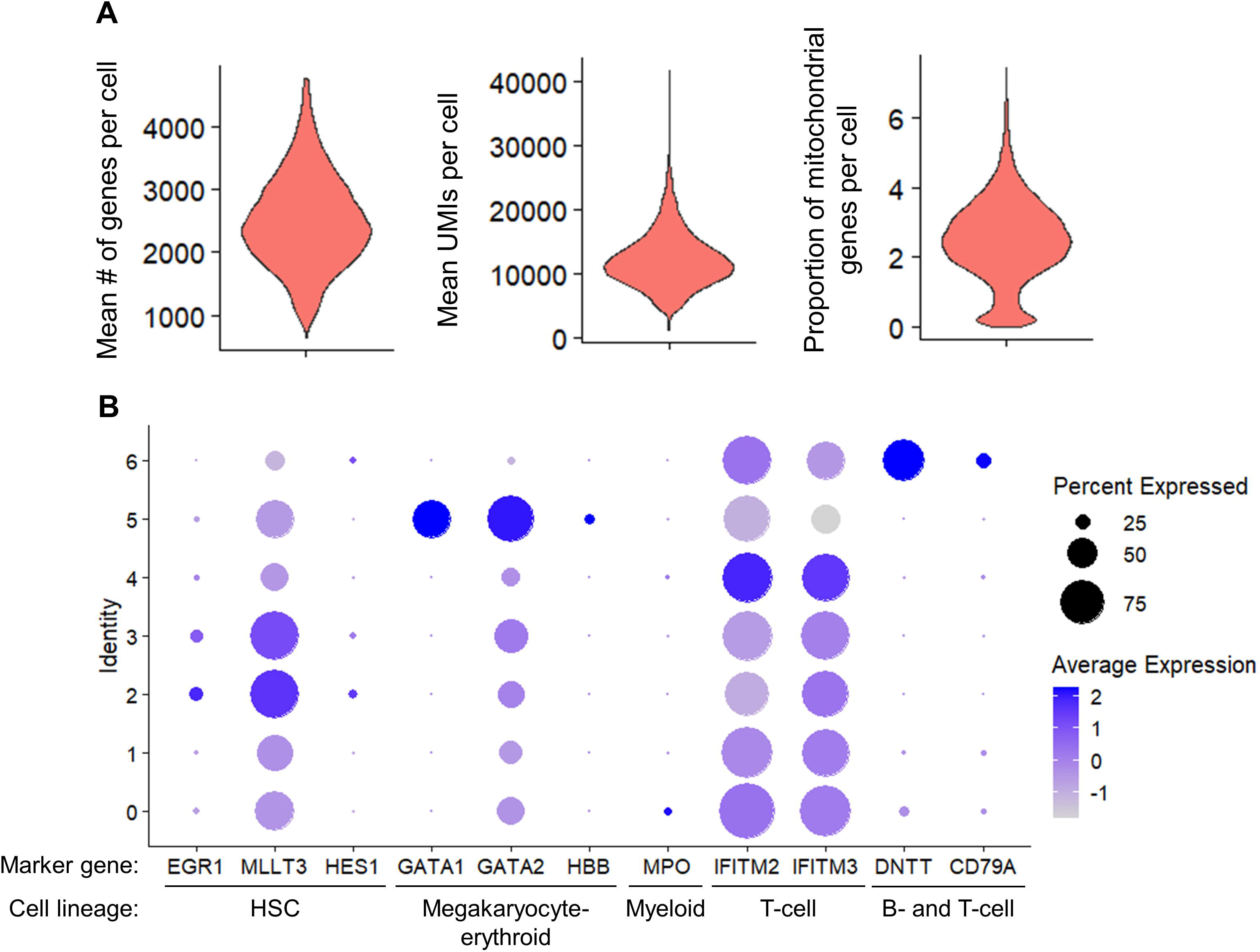

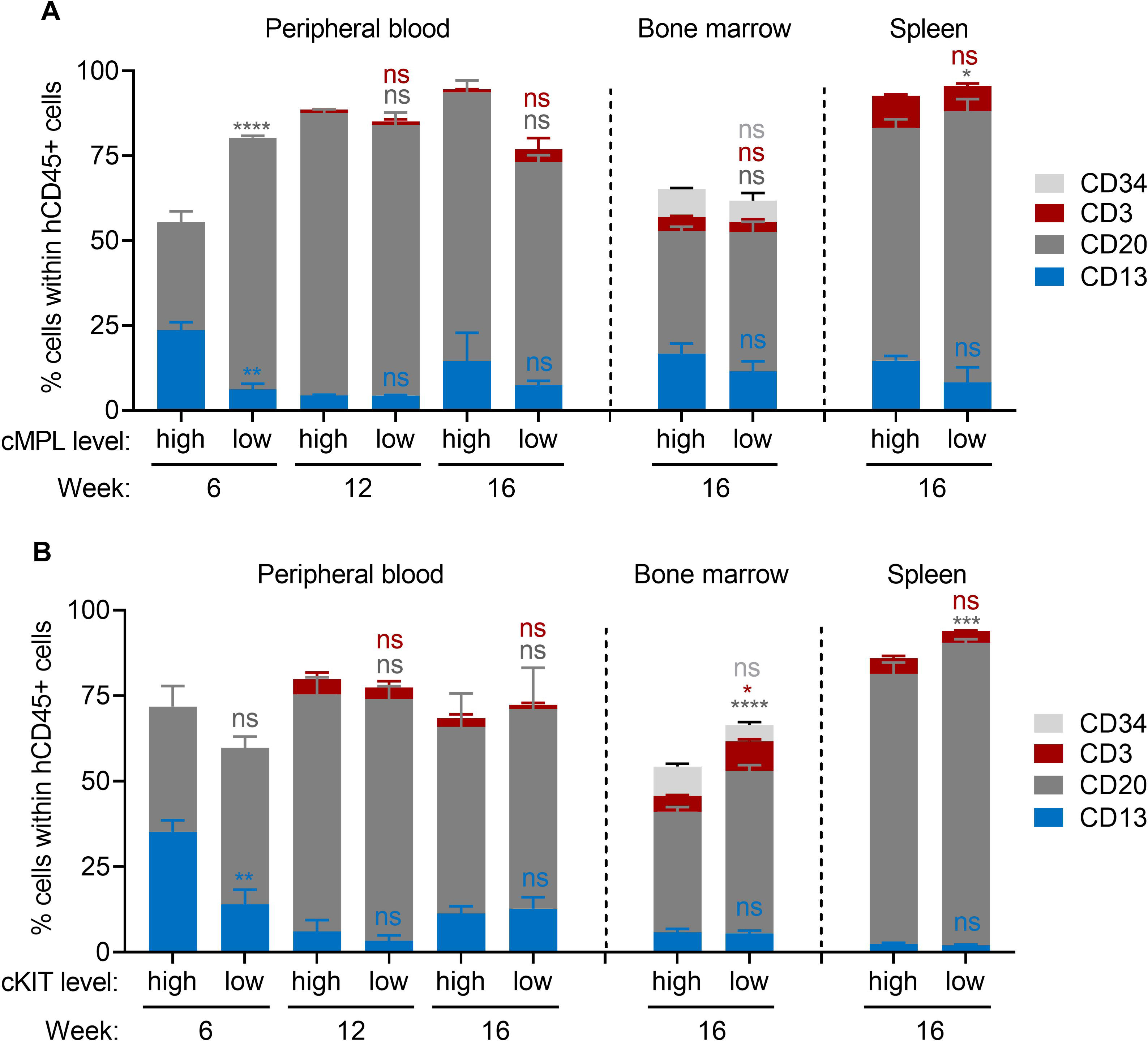

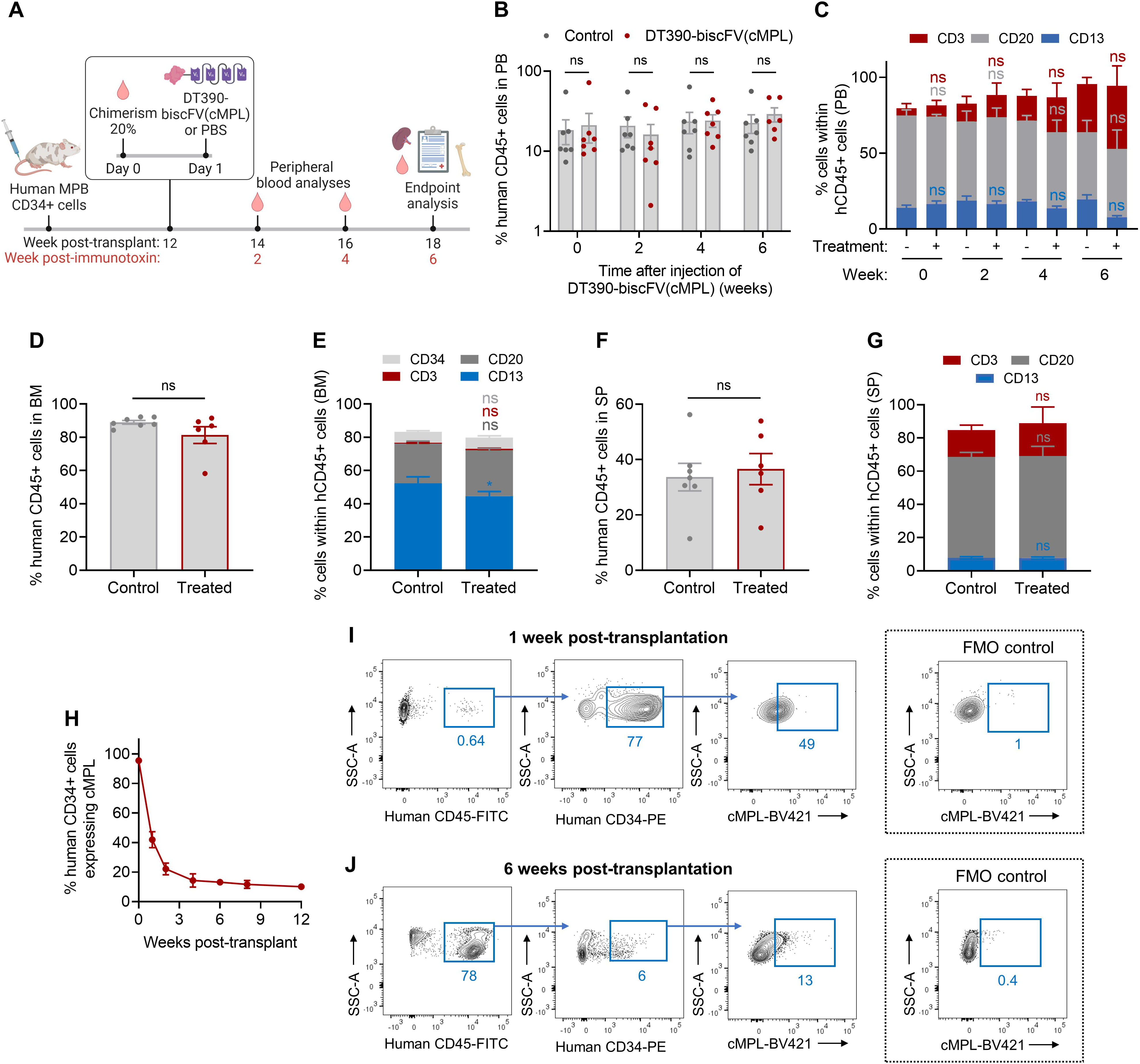

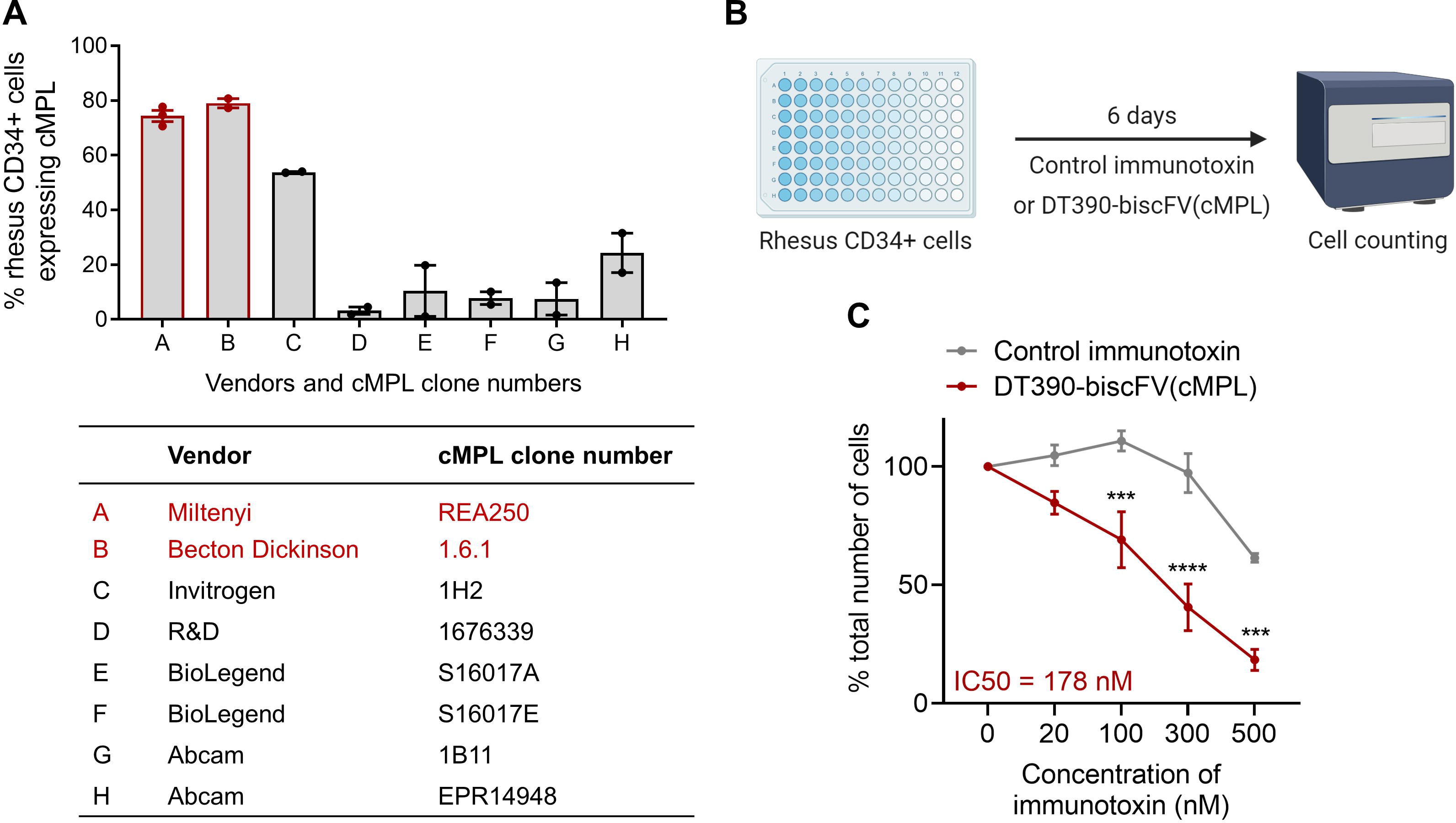

