## Supplemental Information for "cMPL-Based Purification and Depletion of Human Hematopoietic Stem Cells: Implications for Pre-Transplant Conditioning"

Daisuke Araki^1^, Sogun Hong^2^, Nathaniel Linde^2^, Bryan Fisk^3^, Neelam Redekar^3^,

Christi Salisbury-Ruf^1^, Allen Krouse^2^, Theresa Engels^2,4^, Justin Golomb^2,4^, Pradeep Dagur^5^,

Diogo M. Magnani^6^, Zhirui Wang^7^, Andre Larochelle^1^*

**Affiliations:**

(1) Cellular and Molecular Therapeutics Branch, National Heart, Lung and Blood Institute (NHLBI), National Institutes of Health (NIH), Bethesda, MD 20892, USA
(2) Translational Stem Cell Biology Branch, NHLBI, NIH, Bethesda, MD 20892, USA
(3) Integrated Data Science Services (IDSS), National Institutes of Allergy and Infectious Diseases (NIAID), NIH, Bethesda, MD 20892, USA
(4) Priority One Services, Inc., Alexandria, VA 22310, USA
(5) Flow Cytometry Core Facility, NHLBI, NIH, Bethesda, MD 20892, USA
(6) Nonhuman Primate Reagent Resource, University of Massachusetts Medical School, Worcester, MA 01605, USA
(7) Division of Plastic and Reconstructive Surgery, and Division of Transplant Surgery, Department of Surgery, School of Medicine, University of Colorado Denver, Aurora, CO 80045, USA

**Contact information:**

* Corresponding author: Andre Larochelle, M.D. Ph.D., National Heart, Lung and Blood Institute, National Institutes of Health, 9000 Rockville Pike, Bethesda, MD 20892, USA

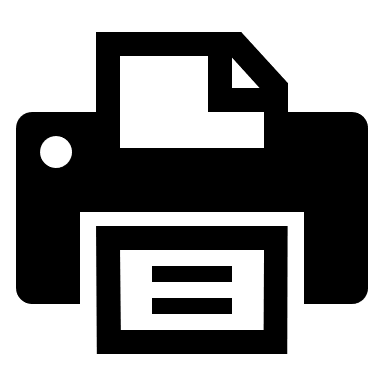

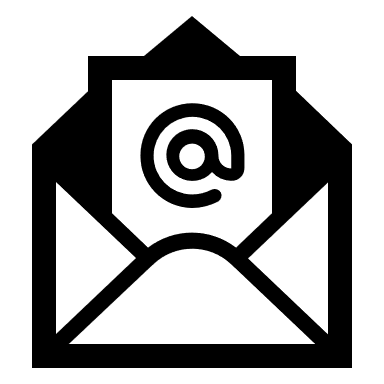

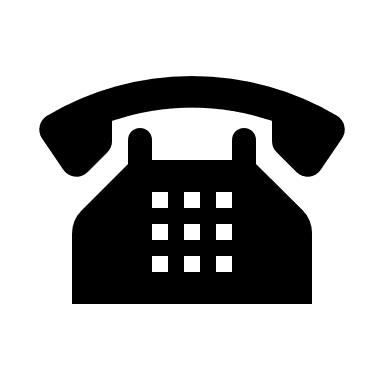

(301) 451-7139	(301) 496-8396

This file includes:

- **Supplemental Figures**
- **Supplemental Figure 1**: Cellular indexing of transcriptomes and epitopes sequencing (CITE-seq) of human mobilized peripheral blood CD34+ cells.
- **Supplemental Figure 2**: Lineage distribution of human CD45+ cells after primary transplantation.
- **Supplemental Figure 3**: Waning cMPL expression after transplantation reduces DT390-biscFV(cMPL) efficacy in depleting engrafted human HSPCs in a xenograft mouse model.
- **Supplemental Figure 4**: Analysis of cross-reactivity of anti-cMPL antibodies between humans and rhesus macaques and the efficacy of DT390-biscFV(cMPL) on rhesus CD34+ cells *in vitro*.

Supplemental Figure 1

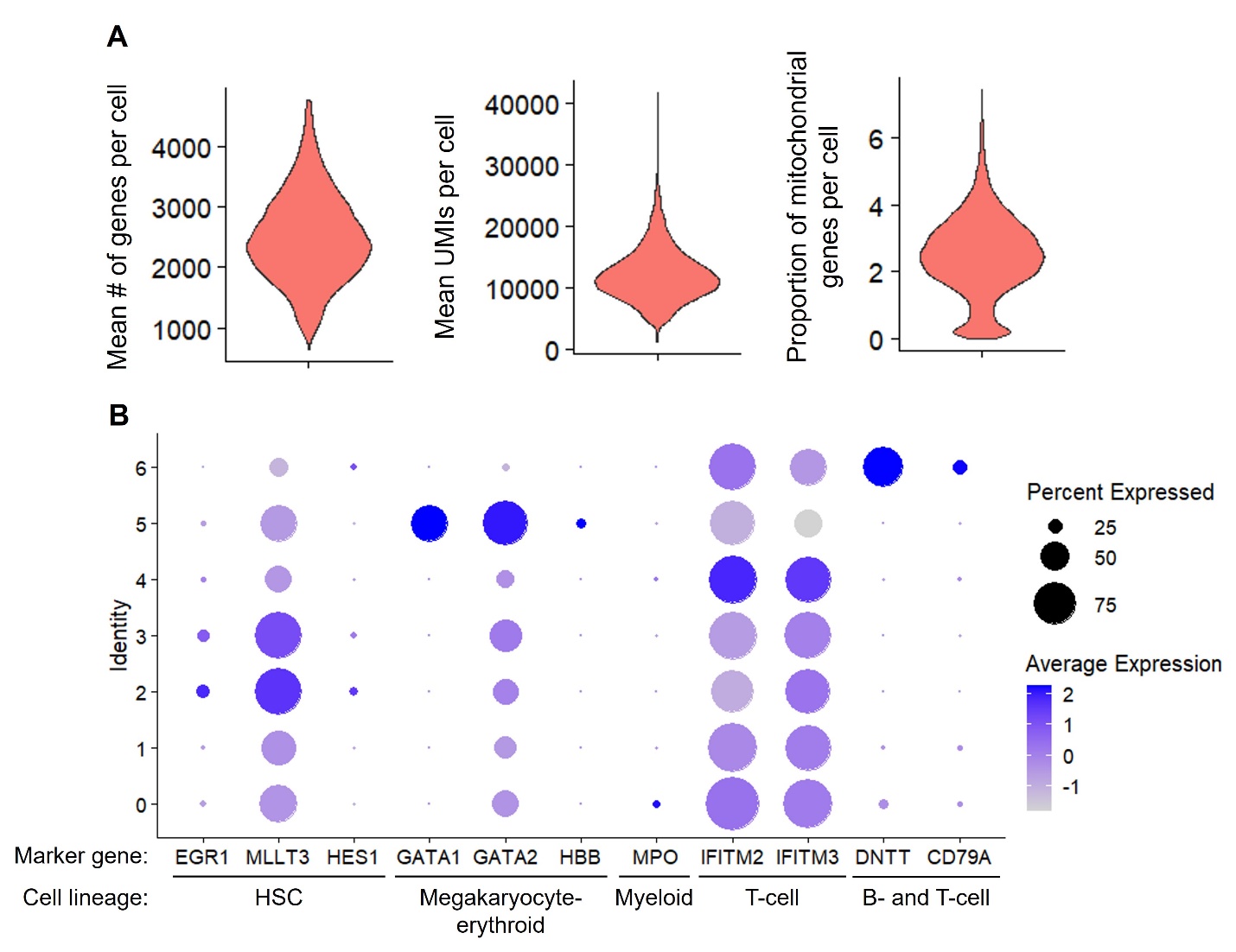

**Supplemental Figure 1 | Cellular indexing of transcriptomes and epitopes sequencing (CITE-seq) of human mobilized peripheral blood CD34+ cells.** (A) Violin plots depicting the distribution of gene counts, transcript counts (Unique Molecular Identifiers, UMIs), and mitochondrial genes per cell after quality control. (B) Dot plot analysis showcasing expression levels of genes associated with hematopoietic stem and progenitor cells across identified cell clusters. Dot size correlates with the proportion of cells expressing each gene, while the color intensity reflects the mean expression level across cluster types. This analysis is associated with findings presented in Figure 1.

Supplemental Figure 2

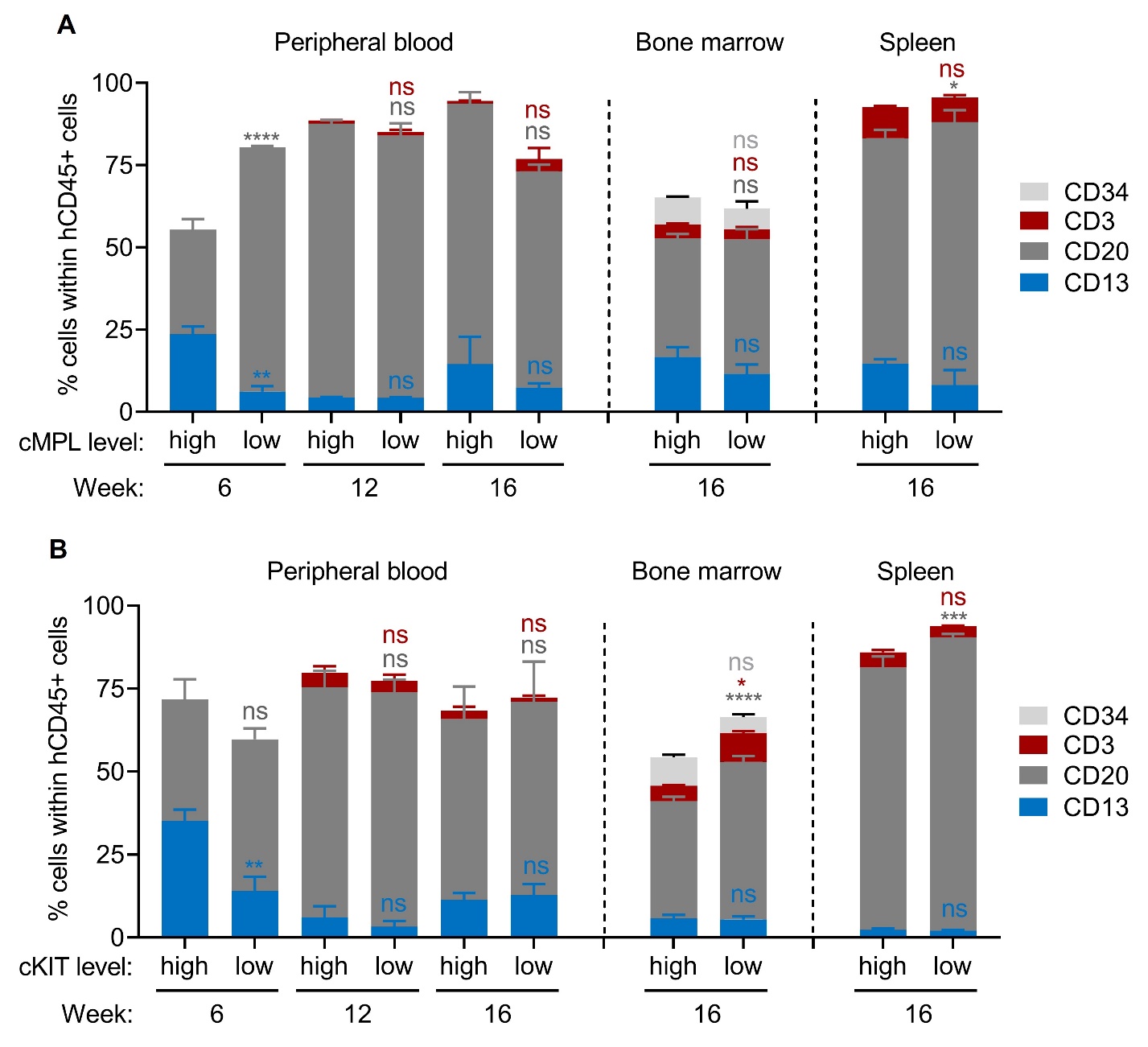

**Supplemental Figure 2 | Lineage distribution of human CD45+ cells after primary transplantation.** (A) Analysis of the lineage distribution within human CD45+ (hCD45+) cells from the peripheral blood (PB), bone marrow (BM), and spleen (SP) following primary transplantation in mice receiving cMPL^high^CD34+ or cMPL^low^CD34+ cells from Figures 2B-D (n = 3-5 mice per group). (B) Analysis of the lineage distribution within hCD45+ cells post-primary transplantation in mice receiving cKIT^high^CD34+ or cKIT^low^CD34+ cells from Figures 2F-H (n = 3-5 mice per group). For the cMPL^low^ group, lineage distribution data was exclusively obtained from mice transplanted with the higher cell dose (2.5 x 10^5^ cells) due to insufficient engraftment in mice transplanted with the lower cell dose (5 x 10^4^ cells). For all other groups, data were collected from mice transplanted with 5 x 10^4^ cells. Datapoints represent the mean ± standard error of the mean (SEM), and significance was evaluated using two-way ANOVA with Sidak's multiple comparisons test. Notations of statistical significance are as follows: ns (not significant), * (p ≤ 0.05), ** (p ≤ 0.01), *** (p ≤ 0.001), and **** (p ≤ 0.0001). This analysis is associated with findings presented in Figure 2.

Supplemental Figure 3
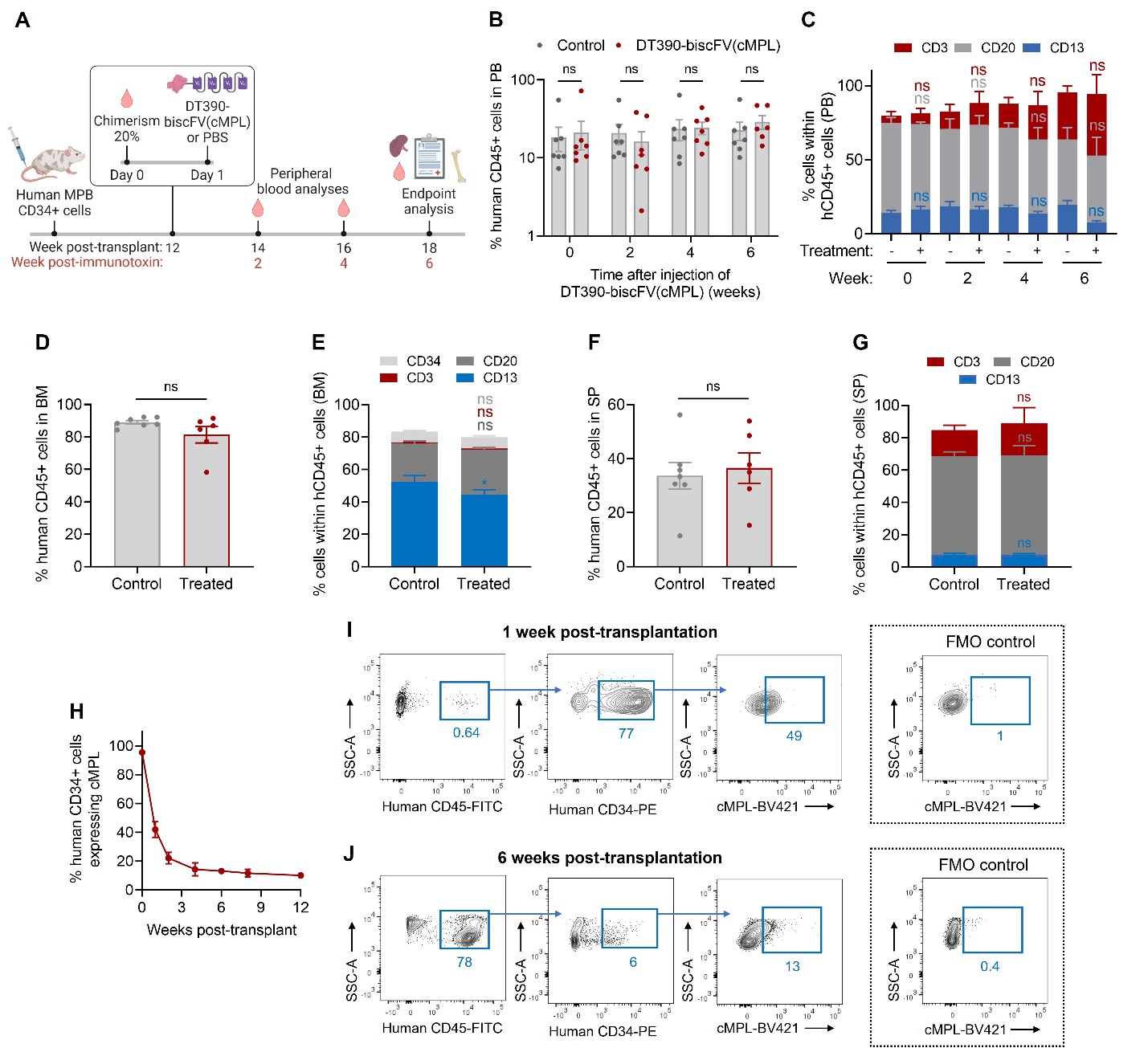

**Supplemental Figure 3 | Waning cMPL expression after transplantation reduces DT390-biscFV(cMPL) efficacy in depleting engrafted human HSPCs in a xenograft mouse model.** (A) Schematic of the experimental procedure: NBSGW mice underwent transplantation with 1x10^6^ human mobilized peripheral blood (MPB) CD34+ cells. At 12 weeks post-transplantation, indicating the establishment of steady-state hematopoiesis, mice received a single dose of either 1.2 mg/kg DT390-biscFv(cMPL) or a PBS control. Human chimerism was monitored in peripheral blood (PB), bone marrow (BM), and spleen (SP). (B, C) Percentage of human CD45+ cells (B) and their lineage distribution (C) in murine PB monitored over a 6-week period following DT390-biscFv(cMPL) administration (n = 6-7 mice per group). (D, E) Percentage of human CD45+ cells (D) and their lineage distribution (E) in murine BM at the endpoint analysis (6 weeks post-injection of DT390-biscFV(cMPL)) (n = 6-7 mice per group). (F, G) Percentage of human CD45+ cells (F) and their lineage distribution (G) in murine SP at the endpoint analysis (n = 6-7 mice per group). (H) Percentage of cMPL+ cells in human engrafted CD34+ cells within murine BM following transplantation with 1x10^6^ human MPB CD34+ cells (n = 4-5 mice per group). (I, J) Representative flow cytometry profiles for human cMPL receptor expression within engrafted human CD34+ cells isolated from murine BM aspirates one week (I) or 6 weeks (J) after transplantation. Fluorescence minus one (FMO) controls set the boundary for positive human cMPL receptor expression at each timepoint. Datapoints represent the mean ± standard error of the mean (SEM), with statistical significance assessed via two-sided unpaired t-tests (panels B, D and F) or two-way ANOVA with Sidak correction (panels C, E and G). Notations of statistical significance are as follows: ns, not significant. This analysis is associated with findings presented in Figure 5.

Supplemental Figure 4
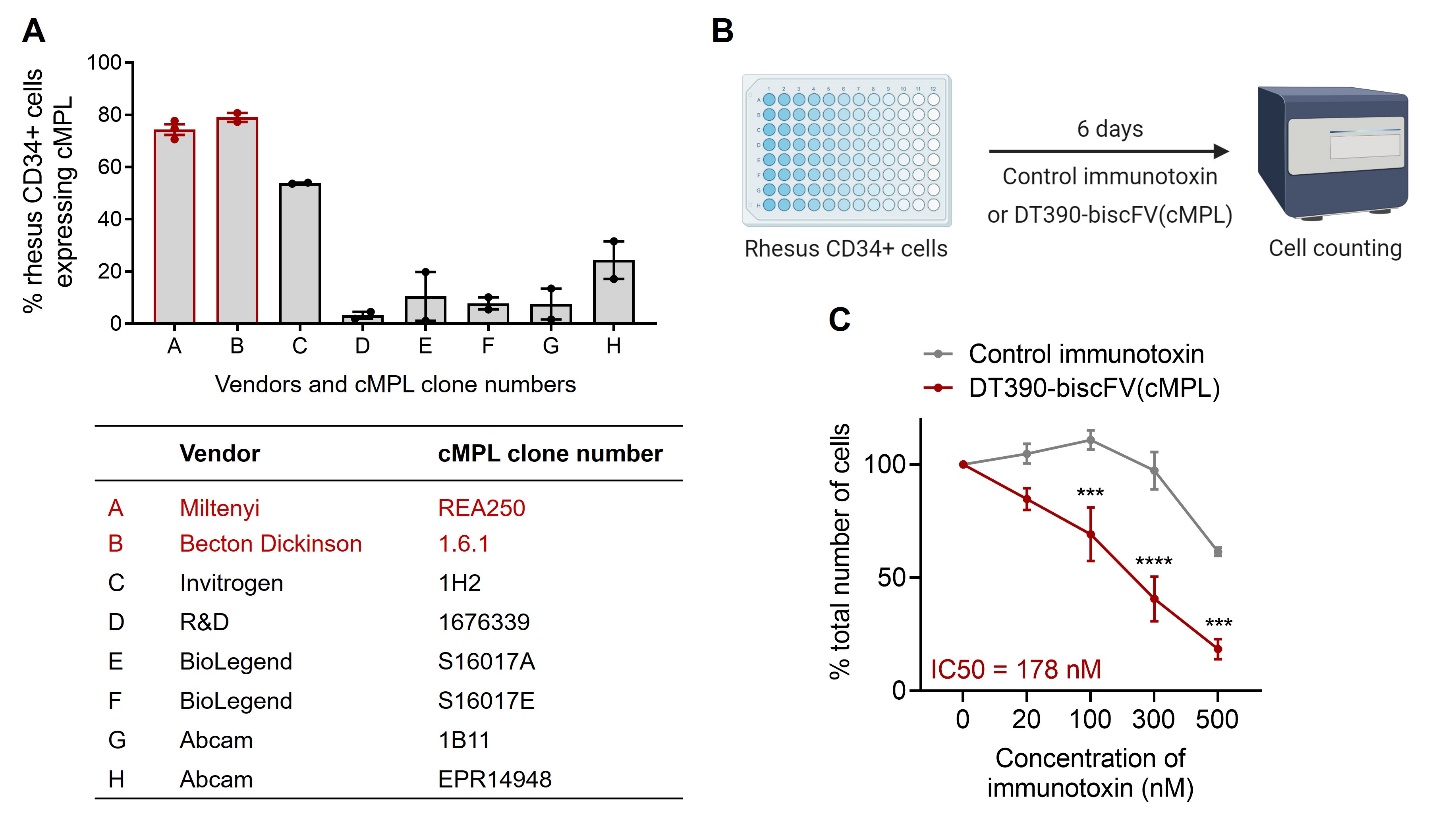

**Supplemental Figure 4 | Analysis of cross-reactivity of anti-cMPL antibodies between humans and rhesus macaques and the efficacy of DT390-biscFV(cMPL) on rhesus CD34+ cells *in vitro*.** (A) Percentage of rhesus CD34+ cells positive for cMPL expression, utilizing anti-cMPL antibody clones from various vendors. Anti-cMPL clones REA250 (Miltenyi Biotec) and 1.6.1 (Becton Dickinson) showed equivalent cross-reactivity with the cMPL receptor on rhesus cells. (B) Schematic of the experimental procedure: *In vitro* cytotoxicity assay was performed using rhesus mobilized peripheral blood (MPB) CD34+ cells. Cells were treated with DT390-biscFV(cMPL) or a non-specific control immunotoxin for 6 days. Cellular growth was assessed by automated counting, with values normalized to untreated control cultures. (C) Response of rhesus MPB CD34+ cells to DT390-biscFV(cMPL) or control immunotoxin after a 6-day culture. Datapoints in panels A and C represent the mean ± standard error of the mean (SEM). In panel C, statistical significance was assessed via two-sided unpaired t-tests. Notations of statistical significance are as follows: *** p ≤ 0.001, **** p ≤ 0.0001). This analysis is associated with findings presented in Figure 6.
